# A molecular map of the human spinal dorsal and ventral horn defines arrangement of neuronal types and glial sex differences

**DOI:** 10.1101/2025.10.31.685953

**Authors:** Katherin A Gabriel, Olivia C Davis, Seph M Palomino, Satoshi Ishishita, Helen Poldsam, Jane M Brandon, Nikhil N Inturi, Hemanth Mydugolam, Ibrahim O Khan, Nethra Selvakumaran, Stephanie Shiers, Muhammad Saad Yousuf, Erin Vines, Peter Horton, Tariq Khan, Anna Cervantes, Geoffrey Funk, Jeffrey C. Reese, Loïs S. Miraucourt, Reza Sharif-Naeini, Jun-Nan Li, Prashant Gupta, Richard A. Slivicki, Ruichen Tao, Robert J. Heuermann, Amol Patwardhan, Gregory Dussor, Eric Meyers, PRECISION Human Pain Network, David Spanswick, Bryan A. Copits, Robert W Gereau, Diana Tavares-Ferreira, Allan-Hermann Pool, Theodore J. Price

## Abstract

The spinal cord is the gateway for somatosensory and nociceptive information to the brain and a key locus for sensory-motor integration. Studies in mice have advanced our understanding of spinal cord circuitry, and transcriptomic studies have begun to characterize the human spinal cord; however, major gaps in knowledge persist. We conducted single-nucleus sequencing of lumbar spinal cord tissue from 11 adult organ donors and annotated spinal cord cell types with high resolution spatial transcriptomics. We identified 34 spatially and transcriptionally defined neuronal classes and detected sex-specific cell types and states across multiple glial populations. Electrophysiological recordings from dorsal horn neurons revealed firing patterns for neuronal subtypes and group I mGluR-dependent plasticity. Our work defines previously unknown aspects of human spinal cord molecular anatomy and physiology.

## Main Text

The spinal cord is the first site of sensory-motor integration in the central nervous system (CNS) with a dorsal horn dedicated to sensory processing, in particular nociceptive information, and a ventral horn that is the final output for motor control from the brain. Many neurological disorders affect the spinal cord. For example, dorsal horn circuitry is frequently linked to chronic pain disorders (*1–3*) and dorsal spinal cord lesions have devastating effects on sensory function (*4, 5*). Motor neuron diseases like Amyotrophic Lateral Sclerosis (ALS) involve degeneration of lower motor neurons in the ventral horn and are often fatal due to a lack of effective treatments(*6*). A recent profiling of human ventral horn motor neurons found that these cells are enriched with genes that contribute to neurodegeneration (*7, 8*), suggesting unique vulnerabilities of these neurons in humans. However, subsequent studies have raised questions about the quality of our current knowledge of human motor neuron transcriptomes (*9*), requiring further validation in larger cohorts. These, and other, previous studies also suggest that there may be important differences in transcriptomes of spinal cord cell types between humans and mice (*8, 10–12*), where most cellular profiling of the spinal cord has been done to date. Indeed, while there are many similarities between mouse and non-human primate (NHP) spinal cord cells, transcriptomic differences in subsets of neurons between mice and NHPs suggest expression differences that may be very important for understanding and treating human diseases that affect or involve the spinal cord (*13*). While other studies have been published profiling all human spinal cord cells in tissues from organ donors, these studies characterized a relatively small subset of cells, used spatial sequencing methods that lack cellular resolution available in current spatial transcriptomic techniques, and did not include sufficient samples from both sexes to examine potential sex differences across all cell types (*8–10, 12, 14*). Although two studies have reported gene expression level differences in some neuronal (*12*) and astrocyte (*10*) populations, sex-specific cell types have remained elusive. A primary goal of our work was to overcome these issues, creating an atlas of human spinal cord neurons and glial cell types with single-cell spatial resolution and a careful examination of potential sex differences across cell types.

There is a strong rationale to design a study to examine sex differences in the human spinal cord. Many chronic pain disorders have sex specific differences in prevalence (*15*) and are thought to involve sensitization of neuronal circuits in the dorsal horn (*1, 2, 16, 17*). Two decades of work has demonstrated that a key cell type driving this plasticity is the primary immune cell of the CNS, microglia (*18, 19*). In neuropathic pain, specific pathways activate microglia in male mice (*20, 21*) leading to brain-derived neurotrophic factor (BDNF) driven modulation of synaptic plasticity that involves both excitatory and inhibitory circuits (*22–24*). Remarkably, this male-specific sexual dimorphism is conserved in human spinal cord (*23*), suggesting that sex differences in dorsal horn microglia may exist in both mice and humans. While there is strong evidence for male-specific pathways in microglia that drive chronic pain, there are other lines of evidence showing that microglia contribute to chronic pain in female rodents, but these underlying mechanisms are only starting to emerge (*25, 26*). Collectively, these experiments indicate that understanding sex differences in human dorsal horn microglia, and other cell types, may be important for engineering new ways to combat chronic pain, but, to date, only one study (*10*) has directly addressed this question in the human spinal cord.

Comparative nervous system transcriptomic studies between other species and humans have revealed major differences including unique cell types that are present in humans (*27–29*), and differences in the composition of receptors, ion channels and peptide neurotransmitters that create challenges in translation of pharmacological findings from other species to humans (*27, 28, 30, 31*). While these differences are now coming into focus in the human dorsal root ganglion (DRG), which projects to the dorsal horn of the spinal cord, and include both unique cell types and important differences in gene expression (*29–31*), far less is known about these potential differences in the dorsal horn. Given that this is the only output pathway for nociceptive information to come from the body to the brain, this represents a critical gap in knowledge that will allow for better translation of spinally targeted therapeutics from preclinical work towards clinical development. Therefore, another major goal of this work was to define the subtypes of neurons in the dorsal horn, gain deeper insights into their transcriptomes, precisely define their spatial location, and compare them to what is currently known in mice using a parallel single-nucleus sequencing method. The data compiled here should serve as a foundation for validation and prioritization of spinally-targeted pain treatments, as well as allowing for mining of new targets that can be investigated for the future treatment of pain, which remains a major unmet medical need (*32, 33*).

## Results

### Quality control of human spinal cord samples from organ donors

Previous studies demonstrate that tissue quality determines transcriptomic features for at least a portion of spinal cord cell types, therefore we sought to determine tissue quality for all samples using strict criteria. Single-nucleus sequencing (snSeq) was performed on 11 lumbar spinal cords (6 females and 5 males) from organ transplant donors (Table 1), using the 10x Genomics platform (Fig. 1A). Three of these donors also underwent FACS sorting using a fluorescently labelled tagged RBFOX3 antibody to enrich for neuronal nuclei (Supplemental Fig.1a). Sequencing quality was high, as confirmed by both technical and biological quality control metrics (Supplemental Table 1) with over 1.2 billion reads, a yield of 161 Gbp, and an efficient 99.45% loading concentration, ensuring deep coverage across nuclei with a median read per nuclei of 8310. At the sample level, nuclei showed excellent transcriptional complexity across all cell types, with median gene counts per nucleus of 2,298 in dorsal horn and 2,279 in ventral horn, and median transcript counts of 3,790 (dorsal) and 3,855 (ventral; Supplemental Table 1). The fraction of reads confidently mapped to the genome was also high, at 85.00% in dorsal horn and 82.20% in ventral horn. Mitochondrial gene expression was low across cell types, with most nuclei exhibiting less than 10% mitochondrial content, consistent with high-purity nuclear isolation (Supplemental Fig. 1B and C). Furthermore, we increased profiling coverage of neurons by enriching for neuronal nuclei with FACS sorting of RBFOX3+/7-AAD double positive nuclei (Fig. 1B and C). Overall, these metrics reflected excellent data quality, that was well-suited for reliable characterization of cellular complexity of the spinal cord.

**Figure 1:**
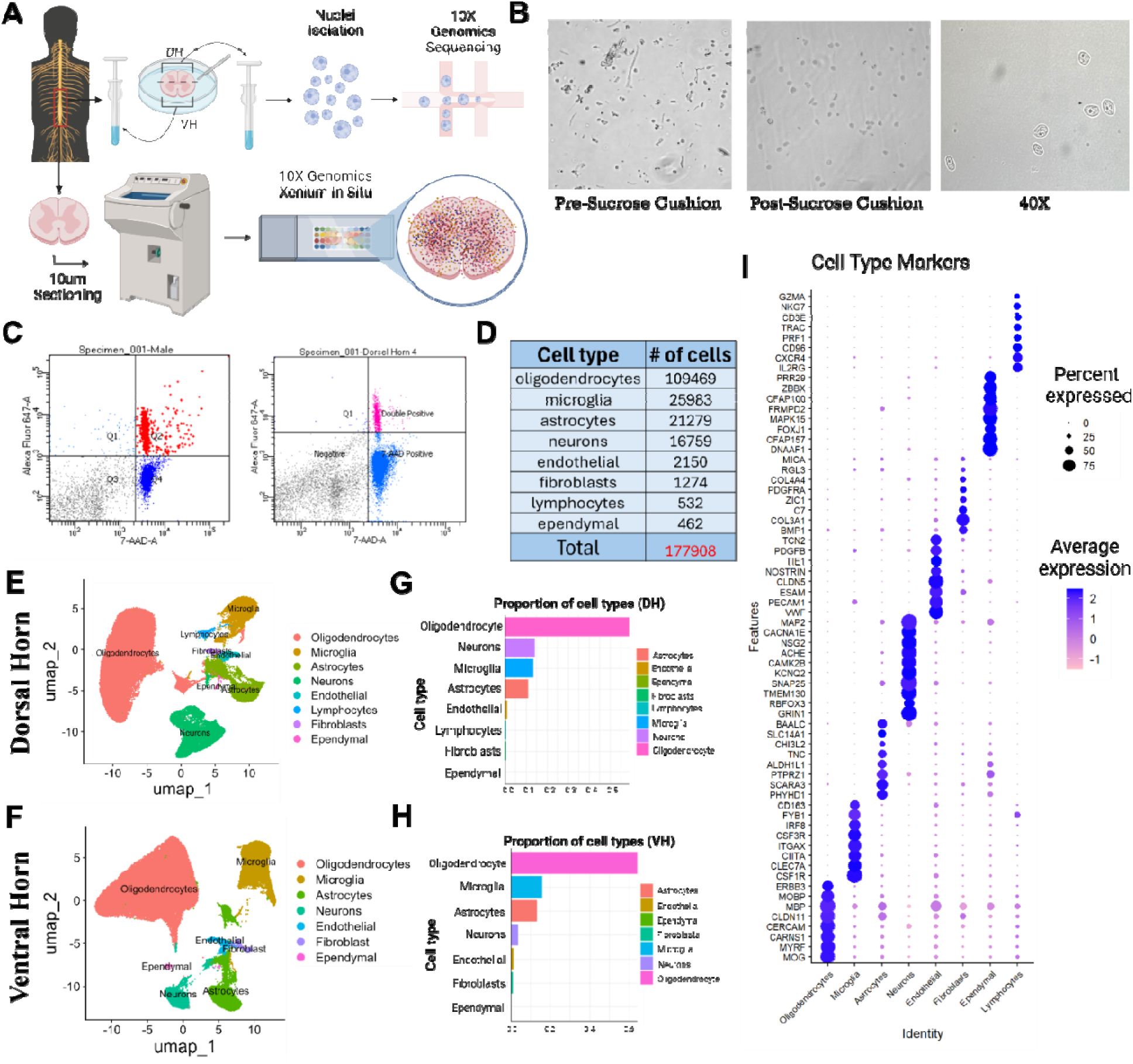
Defining human dorsal horn and ventral horn cell types using single nucleus RNA sequencing. (**A**) Diagram illustrating the two techniques used to generate the data analyzed in this manuscript: single-nucleus RNA sequencing and Xenium spatial transcriptomics. (**B**) Brightfield images of nuclei captured with 20X and 40X objectives on an Olympus IX83 microscope, showing the nuclei before and after sucrose cushion cleanup, demonstrating effective purification and healthy nuclei isolation. (**C**) FACS plots showing double selection of nuclei stained with Alexa-647-RBFOX3 and 7-AAD, used to identify specific neuronal nuclei. (**D**) List of the 8 detected cell types with the corresponding number of cells combined from dorsal horn and ventral horn samples. (**E** to **F**) UMAP plots depicting the 8 different cell types in the dorsal horn (DH) and the 7 cell types detected in the ventral horn (VH), respectively. (**G** to **H**) Bar plots showing the proportions of each cell type in the dorsal horn (DH) and ventral horn (VH). (**I**) Dot plot illustrating the top marker genes expressed across the eight identified cell types.

**Table 1:** Demographic information of donor tissue included in this study.

| **Age** | **Sex** | **COD** | **Ethnicity** | **PMI (Hours)** | **Processing Tissue Used For** | **Medical History** |
| --- | --- | --- | --- | --- | --- | --- |
| 20 | Male | Head trauma / GSW | Black | 2 | Single nucleus sequencing | Surgical history: ACL repair. |
| 23 | Female | Anoxia | White | 2.5 | Single nucleus sequencing | Asthma;<br>Surgical history: Wisdom tooth extraction; Myringotomy; Shoulder arthroscopy (rotator cuff); Surgical repair of right Meniscus |
| 32 | Male | Head trauma / GSW | White | 2.5 | Single nucleus sequencing & Xenium | Approx 20 pack yr smoking history; Excessive drinking ~ 5 nights a week. |
| 36 | Female | Anoxia / OD | White | 3 | Single nucleus sequencing & Xenium | Tox + for THC and Opiates; Anxiety.<br>Surgical history: wisdom tooth extraction. |
| 37 | Female | Head trauma / GSW | Black | 3.5 | Neuronally enriched single nucleus sequencing / Xenium | Insomnia, Major Depressive Disorder. |
| 39 | Male | Anoxia / OD | White | 3 | Neuronally enriched single nucleus sequencing / Xenium | Tox + for Benzo, Cocaine and Opiates; Asthma; 20 pack yr smoking history.<br>Surgical history: Knee/meniscus repair >10 yrs ago. |
| 43 | Male | Head trauma / MVA | Hispanic | 1.5 | Single nucleus sequencing & Xenium | PTSD, Vasectomy |
| 46 | Female | Anoxia | White | 2 | Single nucleus sequencing & Xenium | Tox Screen + Benzos and Marijuana; daily Marijuana use x30 years; Anxiety, Depression, HTN x10 yrs, Hyperlipidemia x2-3 yr; Surgical History: Breast Biopsy, Back surgery, Cervical Scraping, Chest tube placement for lung collapsing. |
| 50 | Female | Head trauma / GSW | White | 2 | Single nucleus sequencing | Depression, Thyroid disease, Asthma, 35+ pack yr smoking history.<br>Surgical History: Hysterectomy, sinus surgery. |
| 56 | Male | Anoxia | White | 2 | Single nucleus sequencing & Xenium | COPD, DM2, HTN, HLD, Asthma; Surgical history: Gallstone, Bilateral Knee replacement (one 10 and one 5 yrs ago), Shoulder surgery >20 yrs ago. |
| 60 | Female | Anoxia | White | 2.5 | Neuronally enriched single nucleus sequencing / Xenium | DM2, Stroke, Cardiac Stents, HTN x 20 yrs, PAD, 20 pack yrs smoking history.<br>Surgical history: Stent in leg, Hysterectomy; |
| 43 | Female | Stroke | White | 2.3 | slice electrophysiology , morphology | Coronary artery disease. Smoking history (0.5-1 pack per day cigarettes for 30 years). Ankle swelling and turned red for the past year. Tubal ligation 8-10 years ago. Drank beer once a month |
| 44 | Female | Stroke | White | 1.7 | slice electrophysiology , morphology | Coronary artery disease. Smoking history (1 pack per day cigarettes for 25 years). Prior fentanyl, amphetamine, benzodiazepine use. Took sleeping medications (Trazadone, Zyprexa, Doxepin). Hard alcohol consumption (10 glasses daily for 25 years). Diagnosed with asthma, didn't use inhaler. |
| 34 | Male | Head Trauma | White | 1.35 | slice electrophysiology , morphology | 2 year cigarette smoking history. Active daily marijuana use (smoking) for 8 years. Daily alcohol consumption (beer). Prior history of smoking methamphetamine (~1/month). Had childhood asthma, took antidepressants & anti-anxiety medications. Lost at least 30 lbs last year. Mother has hypothyroidism. |
| 89 | Female | Stroke | White | 1 | slice electrophysiology , morphology | Drank beer & wine, 1-2 times per week. 2-3 drinks per occasion for the last 50 years. Hypertension |
| 35 | Female | Head Trauma | White | 1.2 | slice electrophysiology , morphology | Daily alcohol consumption (7 shots vodka). Smoked 1 pack per day cigarettes for past 2 years |

\*COD: Cause of Death, GSW: Gunshot wound, MVA: Motor vehicle accident, OD: Overdose, Anoxia: Oxygen deprivation, Stroke: Cerebrovascular accident, PMI: Post-Mortem Interval, ACL: Anterior cruciate ligament, PTSD: Post-traumatic stress disorder, COPD: Chronic obstructive pulmonary disease, DM2: Type 2 diabetes mellitus, HTN: Hypertension, HLD: Hyperlipidemia, PAD: Peripheral artery disease, THC: Tetrahydrocannabinol, Benzo(s): Benzodiazepines, Tox +: Toxicology positive

### Defining the human dorsal horn and ventral horn cell types using single-nucleus RNA sequencing

After quality control, which included removal of nuclei with high mitochondrial read count and exclusion of doublets, we obtained 97,045 nuclei from the dorsal horn and 80,863 nuclei from the ventral horn, totaling 177,908 sequenced nuclei (Fig. 1D). Initially, 30 clusters were identified in the dorsal horn, which were ultimately consolidated into eight distinct cell populations: oligodendrocytes, neurons, microglia, astrocytes, endothelial cells, lymphocytes, fibroblasts, and ependymal cells (Fig. 1E, Supplemental Fig. 1B). In the ventral horn, 27 total clusters were identified and were consolidated to seven distinct cell populations similar to the dorsal horn (Fig. 1F, Supplemental Fig. 1C). A distinct lymphocyte cluster was not detected in the ventral horn unlike the dorsal horn, likely due to their low abundance which may have caused them to fall below detection thresholds during filtering or normalization (Fig. 1F). Regarding cell type proportions, oligodendrocytes were the most abundant cell type in both dorsal and ventral horns, constituting over 50% of the cells (Fig. 1G and H). In the dorsal horn, neurons were the second most abundant cell type at 17%, whereas in the ventral horn, microglia ranked second at 16%, with neurons coming in fourth at 3% after astrocytes at 13% (Fig. 1G and H).

After confirming cell types using known markers from previously published data, we then identified and visualized the top eight highly expressed genes per cluster in a dot plot to highlight cluster-specific marker expression (Fig. 1I). We chose to maintain broad, overarching clusters for initial classification, and subsequently performed sub clustering within each major cell type to explore subtype-level heterogeneity. For example, within the oligodendrocyte population, we observed expression of canonical mature markers such as *MOG, MBP, MOBP, CARNS1*, alongside early-stage progenitor markers like *ERBB3* and *PDGFRA*. Notably, *PDGFRA* was also found as one of the top expressing markers in the fibroblast cluster. We did not detect a clear Schwann cell population, likely due to the extensive removal of dorsal root and grey matter during dissection and pia cleaning, which likely excluded Schwann cells from the nuclei isolation.

### Defining populations of human dorsal horn neurons using single-nucleus RNA sequencing

A total of 11,457 dorsal horn neurons remained following quality control and clustered further into 20 subpopulations (Fig. 2A and B). This is approximately twice the number of neurons that have been sequenced in any single previous human lumbar spinal cord study. More neurons came from female donors than male donors, likely because more tissue from females was processed for neuronally enriched snSeq than males. Despite this, virtually all clusters (19/20) were equally represented in males and females, and no population contained cells from only one sex (Fig. 2C). Several nomenclature systems have been proposed for spinal cord sequencing data (*13, 34*), however we chose to combine multiple aspects from these naming schemes as our data did not fit with an existing spinal cord nomenclature system (*8, 12, 13*). We annotated the neuronal groups with a three-tiered name to denote the main fast neurotransmitter, a differentially expressed gene of historical relevance to rodent literature, and the top differentially expressed gene. In total we found 8 glutamatergic, one cholinergic, one mixed cluster with excitatory and inhibitory nuclei that mostly did not overlap at the single nucleus level, and 10 GABA and / or glycinergic populations defined by expression of genes such as *SLC17A6, CHAT* and *PAX2*, respectively (Fig. 2D).

**Figure 2:**
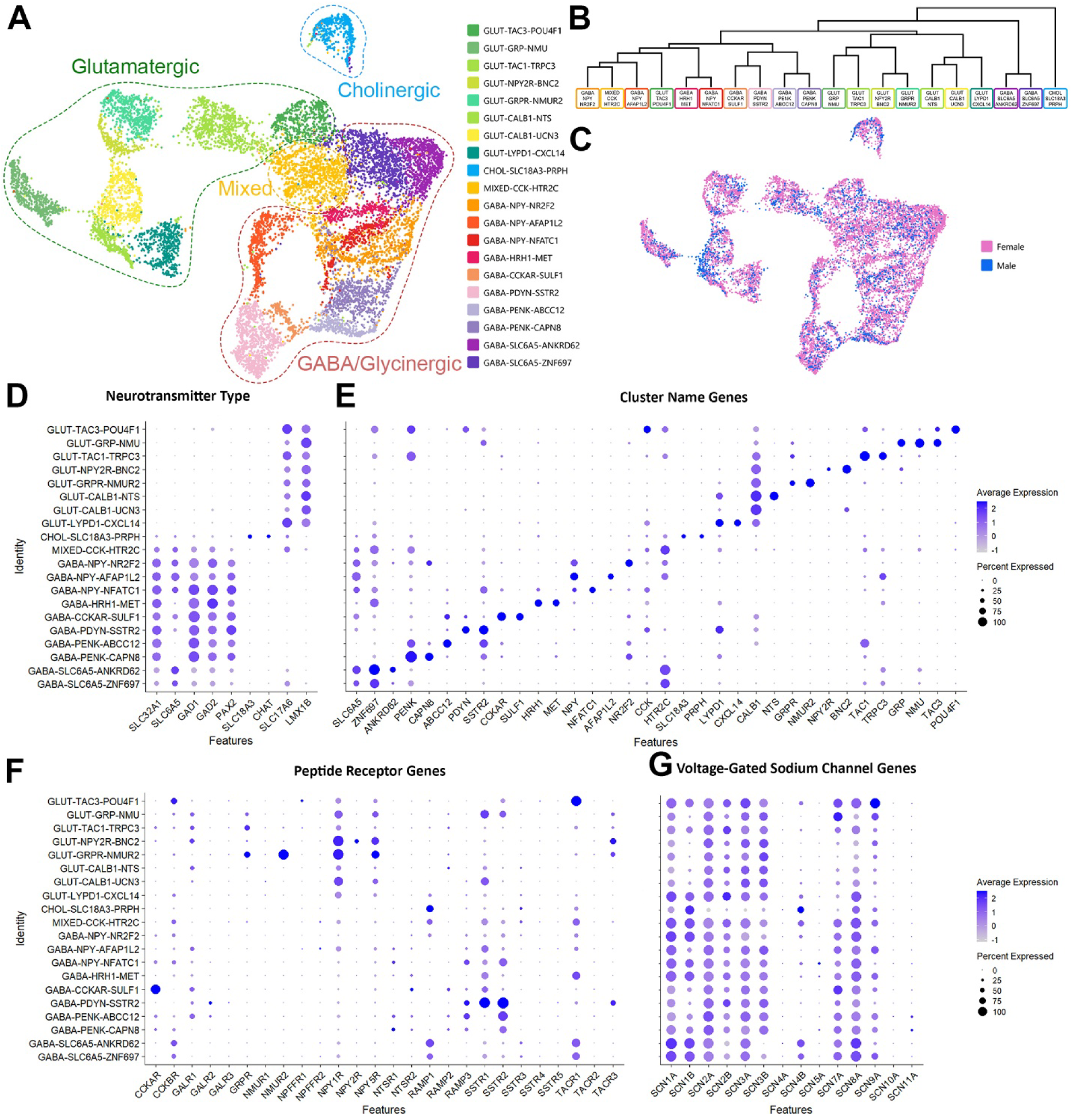
Single nucleus sequencing defines 20 neuronal populations in the human spinal dorsal horn. (**A**) Single nucleus sequencing reveals 20 distinct neuronal populations in the human spinal dorsal horn including 8 glutamatergic, one cholinergic, one mixed glutamatergic and GABAergic, and 10 GABA and/or glycinergic populations; (**B**) hierarchical analysis shows the similarities between clusters at a transcriptomic level; (**C**) cells from females and males are present in all clusters; (**D** to **G**) dotplots showing the expression of fast neurotransmitter markers (**D**), genes used to annotate clusters (**E**), peptide receptor genes (**F**) and voltage-gated sodium channel genes (**G**).

Expression of genes in the second and third tier of cluster names are shown in Fig. 2D. Briefly, several glutamatergic populations contained genes encoding well-defined peptide precursor proteins, GLUT-TAC3-POU4F1, GLUT-GRP-NMU and GLUT-TAC1-TRPC3. Interestingly, some neurons within a separate *GRPR*-expressing cluster (GLUT-GRPR-NMUR2) cluster also express *TAC1* at low levels, suggesting there are two subclusters of *GRPR+* neurons in the human spinal cord distinguished by *TAC1* expression. This finding is in line with a recent preprint that described differential functions to *Grpr+/Tac1+* and *Grpr+/Tac1-* neurons in mice (*35*). We also identified glutamatergic clusters containing the *NPY2R* gene, GLUT-NPY2R-BNC2, and two *CALB1* clusters, defined by co-expression of *NTS* (GLUT-CALB1-NTS) or *UCN3* (GLUT-CALB1-UCN3). The last excitatory cluster, GLUT-LYPD1-CXCL14, contained the *LYPD1* gene.

Consistent with other human spinal cord transcriptomic studies (*8, 12*), we identified one dorsal cholinergic and one dorsal mixed cluster. The cholinergic cluster, CHOL-SLC18A3-PRPH, potentially corresponds to *Chat+* interneurons previously identified in mice (*36*). Whilst these dorsal horn cholinergic neurons have been shown to be inhibitory in rodents, we found very low levels of GABAergic transmission-related genes in this population, leaving uncertainty about their physiological role. Interestingly, the gene encoding the protein peripherin (*<u>PRPH</u>*) was the most differentially expressed gene of this neuronal population. Peripherin expression is thought to be mostly restricted to primary afferent fibers, however, we found peripherin-immunolabelling in cell bodies within the dorsal horn of the human spinal cord alongside central terminal labelling, confirming peripherin expression in human spinal cord neurons (Supplemental Fig. 2A). A mixed dorsal horn cluster containing both excitatory and inhibitory nuclei has not been described in rodent literature but it has been captured in several human and NHP sequencing studies (*8, 9, 12, 13*). We subclustered this population to test whether excitatory and inhibitory neurons would segregate, but this analysis yielded five smaller subclusters, which all still contained a mix of inhibitory and excitatory nuclei (Supplemental Fig. 2B). This suggests that aside from the main neurotransmitter, these cells are highly similar transcriptomically and so were kept as a single cluster for further analysis

We found three inhibitory *NPY+* populations. One marked by expression of *NR2F2* (GABA-NPY-NR2F2), a second marked by *AFAP1L2* (GABA-NPY-AFAP1L2), and the third cluster marked by *NFATC1* co-expression (GABA-NPY-NFATC1). The GABA-NPY-AFAP1L2 population was the only neuronal cluster that showed a sex difference. When comparing the proportion of neurons assigned to this cluster and normalizing for the total number of neurons per donor, the GABA-NPY-AFAP1L2 population represented a significantly larger proportion of neurons in females than males (p = 0.04). However, with 4.6 ± 0.5% and 3.0 ± 1.2% of all neurons in females and males being assigned to the GABA-NPY-AFAP1L2 cluster, respectively, this may not result in a major difference in dorsal horn circuitry and physiology between sexes. More information is needed to draw a solid conclusion.

Two further inhibitory populations were named after differentially expressed receptor genes: the histamine H1 receptor (GABA-HRH1-MET) and the cholecystokinin A receptor (GABA-CCKAR-SULF1). We also found that precursors for endogenous opioids were expressed by a further three clusters of inhibitory interneurons: GABA-PDYN-SSTR2, GABA-PENK-ABBC12 and GABA-PENK-CAPN8. Interestingly, the GABA-PDYN-SSTR2 neurons are potentially analogous to the SST_2a_-expressing dynorphin neurons responsible for somatostatin-induced itch in mice (*37*). Finally, two glycinergic populations containing the gene encoding the glycine transporter 2 (GlyT2, *SLC6A5*), alongside genes involved in GABA neurotransmission, were found to be defined by co-expression of *ANKRD62 (GABA-PENK-ANKRD62)* and ZNF697 (GABA-SLC6A5-ZNF697).

Recent sequencing studies have revealed the expression of peptides among populations of human dorsal root ganglia neurons that project to the dorsal horn of the spinal cord (*29, 31, 38*). Peptides released from these primary afferent central terminals, together with that released by peptidergic interneurons, will activate neuronal circuits expressing peptide receptors. Amongst our dorsal horn populations, we found high expression of several peptide receptor genes including those that bind calcitonin gene-related peptide (CGRP; *RAMP1* and *RAMP3*), neurokinins (*TACR1* and *TACR3*) and somatostatin (*SSTR1* and *SSTR2*), as shown in Fig. 2F.

Voltage gated sodium channels are also of intense interest due to their role in neuronal activity and pain transmission. We found expression of many voltage-gated sodium channels encoding genes in our dorsal horn neuronal clusters (Fig. 2G), and as expected, we did not detect the peripherally restricted channel *SCN10A*, which encodes Na_v_1.8. Expression of other ion channels, ligand-gated receptors and G-protein coupled receptors can be found in Supplemental Fig. 2.

### Defining populations of human ventral horn neurons using single-nucleus RNA sequencing

As expected, fewer neurons were identified by snSeq of the ventral horn compared to the dorsal horn, but these neurons could be split into 14 subclusters (Fig. 3A and B). Consistent with the dorsal horn, more of these neurons came from female samples than male samples. However, all clusters were equally represented in males and females when normalized to the total number of neurons per donor with no statistical differences, and no cluster had cells from only one sex (Fig. 3C).

**Figure 3:**
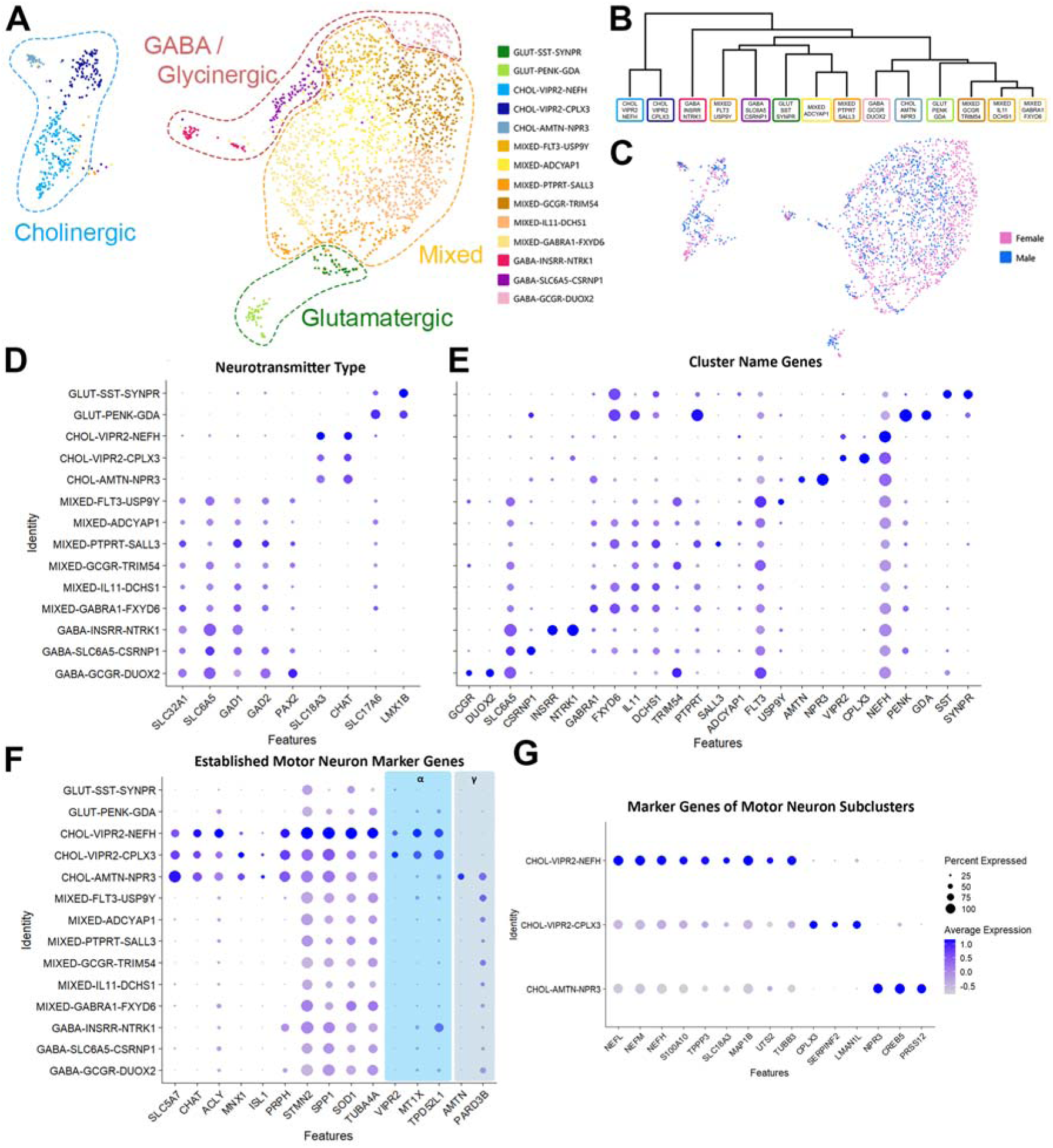
Single nucleus sequencing defines 14 neuronal populations in the human spinal ventral horn. (**A**) Single nucleus sequencing reveals 14 distinct neuronal populations in the human spinal ventral horn with 2 glutamatergic, 3 cholinergic motor neuron, 6 mixed glutamatergic and GABAergic, and 3 GABA and/or glycinergic populations; (**B**) hierarchical analysis shows the similarities between clusters at a transcriptomic level; (**C**) cells from females and males are present in all clusters; (**D** to **G**) dotplots showing the expression of fast neurotransmitter markers (**D**), genes used to annotate clusters (**E**), established marker genes of motor neurons, including alpha and gamma-specific genes and distinguishing marker genes of the three motor neuron clusters defined in this study (**G**).

The ventral horn clusters were named using the same system described above, resulting in 2 glutamatergic, 3 cholinergic, 6 mixed with both excitatory and inhibitory nuclei, and 3 GABA and/or glycinergic populations (Fig. 3D). Potential genes of interest expressed by each population formed the second tier of the name, and the most differentially expressed gene was used to create the third and final tier (Fig. 3E). Dot plots demonstrating expression of other genes including those encoding ion channels and receptors can be found in Supplemental Fig. 3.

Only 2 of the 14 ventral horn populations identified in the ventral horn using snSeq were established as glutamatergic based on differentially expressed genes. While *SST* and *PENK* were widely expressed across several dorsal horn populations, these genes each defined a single excitatory cluster in the ventral horn: GLUT-SST-SYPR and GLUT-PENK-GDA.

We also found six clusters in the ventral horn with excitatory and inhibitory nuclei, all of which had relatively similar transcriptomic signatures. Again, we grouped the cells in these six mixed populations together and re-clustered them in an attempt to gain more distinction between excitatory and inhibitory neurons, but this only yielded seven subclusters (Fig. S3A), all of which still contained excitatory and inhibitory nuclei. This suggests that more extensive cellular profiling may be required to resolve stable transcriptomic identities within these mixed clusters.

As sub clustering did not offer any distinction between excitatory and inhibitory populations, we decided to keep them as the six mixed clusters for further analysis. Of these six transcriptomic classes, one population (MIXED-ADCYAP1) was defined by a single differentially expressed gene, *ADCYAP1*, which encodes the peptide PACAP, although the remaining mixed clusters were defined by multiple differentially expressed genes. These include MIXED-PTPRT-SALL3, MIXED-GCGR-TRIM54, MIXED-IL11-DCHS1 and MIXED-GABRA1-FXYD6. The latter population contained cells with a high expression of *EPHA4*, a marker of ipsilaterally projecting interneurons in mice that are involved in central pattern generator circuits and can directly activate motor neurons (*39*). The last mixed cluster, MIXED-FLT3-USP9Y, had a Y-linked gene as the most differentially expressed gene, which formed the third tier of the cluster annotation. However, this population was represented equally in females and males and is not male-specific.

Three ventral inhibitory clusters were found including GABA-INSRR-NTRK1 and GABA-GCGR-DUOX2, although only one contained differentially expressed genes related to glycinergic as well as GABAergic transmission (GABA-SLC6A5-CSRNP1). This glycinergic cluster may also include human Renshaw-like cells, as some neurons in this population also express *CALB1* and / or *PVALB*, which are commonly co-expressed markers of V1-derived Renshaw cells in mice (*40*).

All three ventral cholinergic groups in this study were identified as motor neurons based on the expression of *SLC18A3* (vAChT), *MNX1*, *ISL1* and *CHAT* (Fig. 3F). In total, there were 333 neurons in these motor neuron clusters, which accounted for approximately 0.2% of all cells in this study (333 / 177,634). This is consistent with previous estimates of the incidence of motor neurons in human spinal cord (*9*). Two of the motor neuron clusters had a high expression of *VIPR2* (CHOL-VIPR2-NEFH and CHOL-VIPR2-CPLX3), together with other markers of alpha motor neurons and therefore likely represent this subtype of motor neuron in humans (*41*). *CPLX3* was the most differentially expressed gene in one of these *VIPR2-*expressing groups, in line with a recent preprint that also found *CPLX3* was a specific marker of human alpha motor neurons and was downregulated in the spinal cord of patients with ALS (*42*). The third motor neuron cluster (CHOL-AMTN-NPR3) contained high levels of *AMTN*, *CREB5* and *PARD3B,* which have been shown to be markers of gamma motor neurons in human (*9*). Together, this suggests motor neurons in our study were sequenced at a sufficient depth to separate out the two types of motor neuron, which has been an area of contention in previous human spinal cord transcriptomic studies (*8, 9*). Gautier et al., (2023) also showed that neuronal “debris” can contaminate human ventral horn sequencing studies, leading to a low-quality motor neuron population that exhibits significantly lower gene counts and other quality control variables in comparison to the remaining neurons. We undertook several rounds of doublet removal during quality control to remove any debris-enriched populations with low gene counts, however, we were still concerned the CHOL-VIPR2-NEFH population could be contaminated with axonal debris due to the high expression of cytoskeletal genes. We found that the unique genes and UMIs of all three motor neuron groups were comparable to the remaining ventral horn groups, suggesting no overt contamination of low-quality debris in any cholinergic population (Supplemental Fig. 3B). We also found the average expression of NEFH was ∼8-fold higher in neurons in the CHOL-VIPR2-NEFH cluster than the CHOL-VIPR2-CPLX3 population and ∼13-fold higher in the CHOL-AMTN-NPR3 cluster (Fig. 3G), despite being expressed in virtually all neurons in all three populations (>98%). This enrichment of cytoskeletal genes and quality of the nuclei within this cluster suggests CHOL-VIPR2-NEFH is a third population of motor neurons, which is distinct from other alpha motor neurons expressing *CPLX3* and gamma motor neurons defined by *ATMN-*expression (Fig. 3G).

### Spatial architecture of the human spinal cord revealed using Xenium spatial transcriptomics

Utilizing the top expressing markers of each major cell type from snSeq, we developed a 480-custom gene Xenium spatial transcriptomics panel to determine the localization of these cells within the architecture of the spinal cord (see Supplemental Table 2 for a list of genes included). Here, we show a representative output from two spinal cord samples, one from a 36-year-old female and the other from a 32-year-old male, processed using the 10x segmentation kit (Fig. 4A and B). Although the Xenium segmentation kit notably improved the overall segmentation accuracy, it remained challenging to reliably detect larger neurons including motor neurons. To overcome this limitation, we manually segmented all neuronal cells using the composite images generated in the Xenium processing stages to generate a reference map. This was then integrated with the Xenium output data for each sample, thereby enhancing the accuracy of neuronal detection used for further analysis. Consequently, for each of the 8 samples, we removed empty cells, normalized gene expression, reduced dimensionality, clustered and then plotted the cells (Fig. 4C). We generated and named clusters from each of the 8 samples after verifying with the top marker genes (Supplemental Fig. 4). We then moved on to integrating all the donors using SCT-based normalization, aligned shared cell types across batches, and performed clustering and visualization on the merged data (Fig. 4D). We obtained a total of 39 clusters and looked at the top expressing genes for the major cell types generated by the snSeq to determine the identity of each individual cluster (Fig. 4E and F). After normalization and integration, we were left with 704,685 total cells, and 16,568 neurons collectively from all of the 8 samples (Fig. 4E). We then validated the top expressing genes identified from the snSeq data using the Xenium integrated clusters as visualized in the dot plot (Fig. 4G).

**Figure 4:**
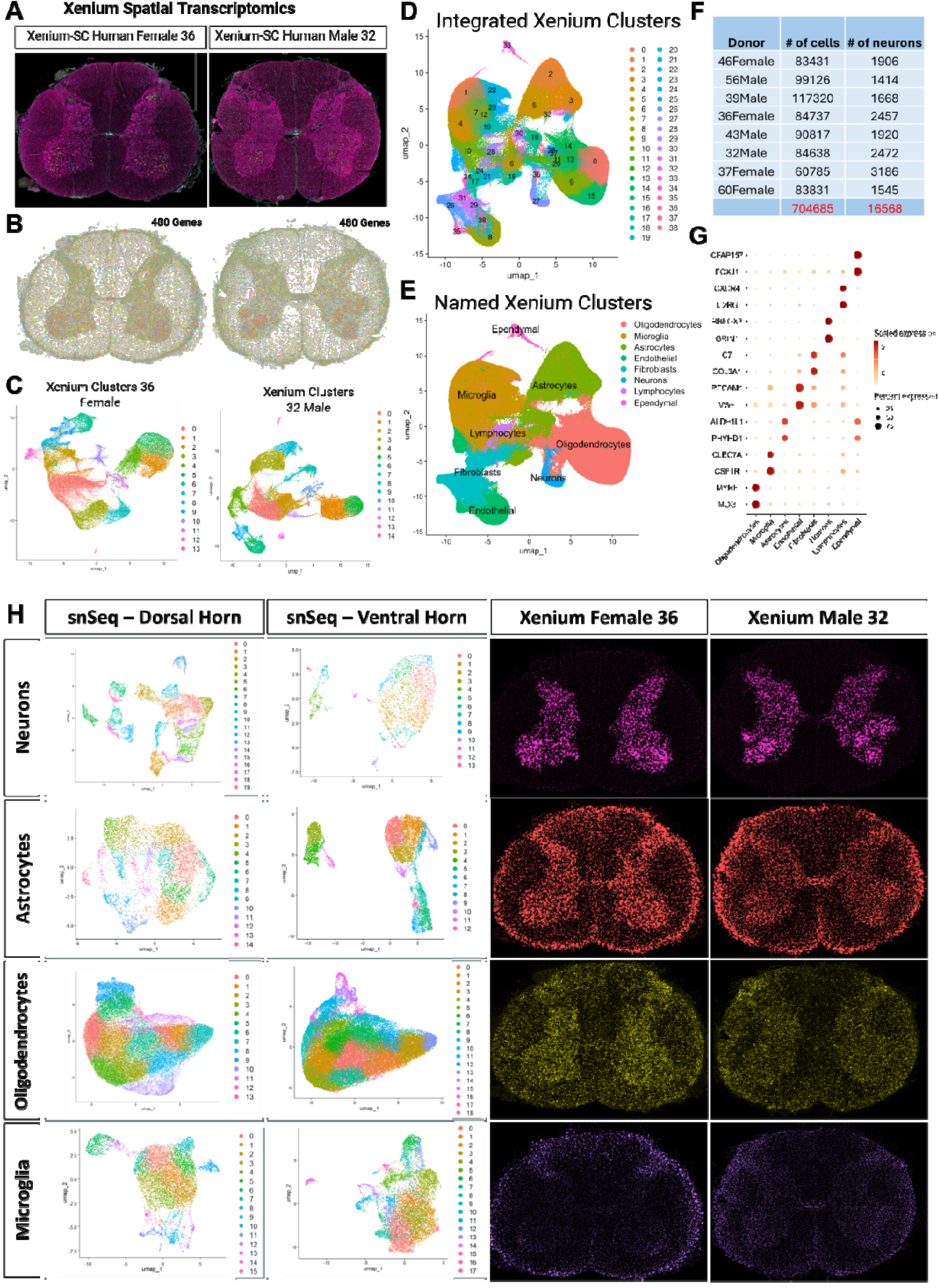
Spatial architecture of the human spinal cord revealed using Xenium spatial transcriptomics. (**A)** representative output image of xenium spatial transcriptomics using the cell segmentation kit in xenium explorer of one male and one female sample. (**B**) representative image of all 480 genes of interest mapped onto the spinal cord section of one male and one female sample. (**C**) UMAP plots depicting all of the different cell types detected in each representative male and female samples. (**D**) UMAP of all 8 integrated xenium samples. (**E)** UMAP plot of all 8 integrated xenium samples representing all 8 cell types detected: oligodendrocytes, neurons, microglia, astrocytes, lymphocytes, endothelial cells, ependymal cells and fibroblasts. (**F)** List of total number of cells and total number of neurons detected in each of the 8 xenium samples. (**G)** Dotplot depicting the top 2 genes of interest used to categorize each cluster, demonstrating the top expressing markers generated in the single nuclei sequencing also represent the top expressing markers in the xenium spatial transcriptomics. (**H**) subclustered UMAP plots of the top 4 cell types using the single nuclei sequencing both in the dorsal horn and ventral horn, and it’s corresponding spatial architecture using the xenium spatial transcriptomics of a representative male and female samples.

Next, we showcased the subclusters of the four most abundant cell types (neurons, astrocytes, oligodendrocytes, and microglia) analysed separately in the dorsal and ventral horns (Fig. 4H). Both neurons and astrocytes exhibited a greater number of clusters in the dorsal horn (20 and 15 clusters, respectively) compared to the ventral horn (14 and 13 clusters, respectively), reflecting the increased complexity of sensory integration within the dorsal horn and highlighting the likely role of astrocytes in synaptic modulation. In contrast, oligodendrocytes and microglia showed more clusters in the ventral horn (19 and 18 clusters, respectively) than in the dorsal horn (14 and 16 clusters, respectively). Given the high metabolic demands of motor neurons, the presence of specialized oligodendrocytes for metabolic support, along with enhanced immune surveillance by microglia, this finding is consistent with the functional requirements of ventral horn motor neurons (*43*). To further elucidate the spatial organization of these cell types at single-cell resolution, we utilized Xenium transcriptomics to map the top-expressing markers of all four cell types in representative male and female donors, as illustrated in Figure 4H.

### Spatial characterization of human dorsal horn neuronal populations

To determine the laminar distribution of neuron clusters identified in the snSeq data, cells with at least 3 transcripts of each marker gene per neuronal population and RBFOX3 (to exclude non-neuronal cells) were visualized using Xenium Explorer (dot plots showing the expression of each dorsal horn population marker gene can be found in Supplemental Fig. 2). These images were overlaid onto a lumbar spinal cord atlas adapted from (*44*) and manually plotted in space (all donors overlaid in Fig. 5A, see Supplemental Fig. 5A for individual plots). All clusters contained cells from both males and females with no clear sexual dimorphism in the spatial organization of neuron types (see Supplemental Fig. 5B for plots divided by sex).

**Figure 5:**
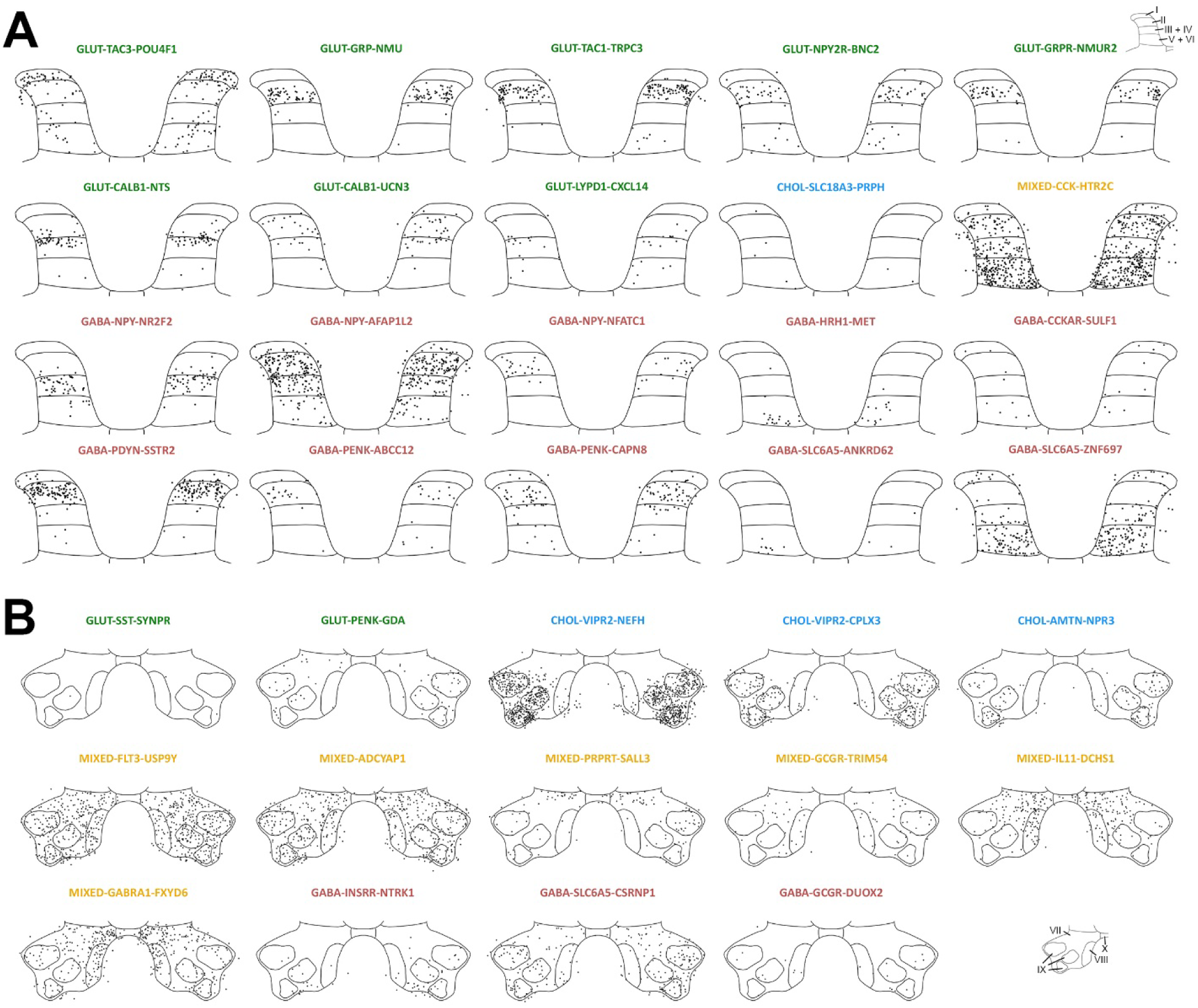
Laminar distribution of neuronal populations in the human spinal dorsal and ventral horn. (**A**) Spatial distributions of 20 dorsal horn neuronal clusters identified using Xenium technology, plotted onto a dorsal horn atlas for visualization and comparison between donors. (**B**) Spatial distributions of 14 ventral horn neuronal clusters identified using Xenium technology, plotted onto a ventral horn atlas for visualization and comparison between donors. Both dorsal and ventral horn maps are overlays of the outputs for 8 donors, (not to scale, each dot represents a single cell).

The GLUT-TAC3-POU4F1 population occupied lamina I as well as the deeper laminae. Whilst this does not match *Tac2* (the orthologous mouse gene encoding neurokinin B) expression in rodents (*45*), it does resemble the *Pou4f1* expression pattern observed embryonically in mice (*46*). Several other glutamatergic populations in our study, GLUT-GRP-NMU, GLUT-TAC1-TRPC3 and GLUT-GRPR-NMUR2, contained neurons largely restricted to lamina II, consistent with *Grp*, *Tac1* and *Grpr* expression patterns in the rodent (*47–49*). The GLUT-NPY2R-BNC2 subcluster mostly had cells in lamina II, with some scattered throughout laminae V-VI. Of the two *CALB1* clusters, one was located only in lamina III (GLUT-CALB1-NTS) and the other was spread throughout laminae II-IV (GLUT-CALB1-UCN3). This peptidergic expression appears to be expanded in humans, as *Calb1*, *Nts* and *Ucn3* have been shown to be restricted to the superficial laminae in rodents (*50–52*). The last excitatory cluster, GLUT-LYPD1-CXCL14, contained cells from lamina II_inner_ to lamina VI. The rodent ortholog of this gene, *Lypd1,* is a marker of spinoparabrachial neurons in the mouse and these cells often co-express *Tacr1*(*53*). A small subcluster of the neurons within the GLUT-LYPD1-CXCL14 cluster also co-express *TACR1*, suggesting these markers of spinoparabrachial neurons may be conserved across species. However, *TACR1* is more highly expressed in the GLUT-TAC3-POU4F1 population and as cells in that cluster are also localized to lamina I and deeper dorsal horn laminae, this population may also contain ascending projection neurons. The GLUT-TAC3-POU4F1 cluster also appears to contain some cells in the white matter adjacent to the lateral edge of the grey matter, possibly revealing the lateral spinal nucleus (LSN) in humans. In rodents, projection neurons that form part of the anterolateral system are found in this nucleus (*54*), although it is unclear whether the LSN is present in human spinal cord as these cells were not present in tissue sections from all donors.

Neurons in the dorsal cholinergic cluster were sparsely scattered throughout the deeper laminae (CHOL-SLC18A3-PRPH). This small population likely reflects a corresponding cholinergic interneuron group observed in rodents, with a previous study identifying only 24 GFP+ neurons in the dorsal horn per spinal segment in a *Chat*^eGFP^ transgenic mouse line (*36*).

The mixed dorsal horn cluster, MIXED-CCK-HTR2C, contained neurons scattered throughout the dorsal horn with a particularly dense band of neurons in laminae V-VI. This is consistent with other human spinal cord atlases that found a mixed deep dorsal horn cluster using Visium technology (*8, 12*), although as discussed previously, a corresponding population has not been described in rodent literature and the potential role in sensory circuitry is unknown.

Of the three inhibitory *NPY–*containing populations, GABA-NPY-NR2F2 was located in laminae III and IV, GABA-NPY-AFAP1L2 had neurons throughout laminae II to IV, and the third cluster GABA-NPY-NFATC1 was restricted to lamina II. Three subclusters of inhibitory *Npy*-expressing neurons were also described in a mouse dorsal horn transcriptomic study, and whilst they were not defined by the same marker genes as this study, the spatial organization is similar with cells scattered throughout laminae I – IV (*53*). The GABA-HRH1-MET subcluster was restricted to the deeper laminae, while the GABA-CCKAR-SULF1 population had cells in lamina I as well as the deep dorsal horn. GABA-PDYN-SSTR2 neurons were restricted to lamina II with a similar laminar distribution to the SST_2a_-expressing dynorphin neurons in mice (*37*). Two subclusters with *PENK*+ neurons, GABA-PENK-ABBC12 and GABA-PENK-CAPN8, were mainly restricted to superficial laminae with only a few cells in the deeper laminae, consistent with the distribution of GFP+ cells in the spinal cord of *Penk*^eGFP^ BAC transgenic mice(*55*). Finally, the smaller glycine-containing population, *GABA-SLC6A5-ANKRD63, was* mainly found in laminae V and VI; while the other glycinergic population, GABA-SLC6A5-ZNF697, also had cells in lamina I as well as the deeper dorsal horn. These neurons appear to be distributed in the dorsal horn in a similar pattern to tdTomato-expressing cells in spinal cord sections of a GlyT2^tdTomato^ transgenic mouse line (*56*).

### Spatial characterization of ventral horn interneurons and motor neurons

The 14 subclusters of ventral horn neurons were also plotted into space using 10X Xenium technology and showed less distinct spatial organization than the dorsal horn neurons (Fig. 5B) for ventral horn plots with all donors overlaid and Supplemental Fig. 6A and B for a breakdown for each donor and when grouped by sex).

Neurons in the GLUT-SST-SYNPR cluster were scattered sporadically throughout the ventral horn, reflecting the sparse pattern of a small number of somatostatin-immunoreactive neurons in the ventral horn of the rat (*57*). Cells belonging to the other ventral glutamatergic population, GLUT-PENK-GDA, were also found throughout the ventral horn, excluding lamina VIII.

All three motor neuron populations were restricted to lamina IX nuclei, as expected, but there were no obvious differences in the organization between these groups. We found many examples of large motor neuron nuclei in lamina XI that expressed markers of the CHOL-VIPR2-NEFH cluster, cholinergic genes (such as CHAT) but lacked markers of the other two motor neuron populations (Supplemental Fig. 6C). This further supports the notion that the CHOL-VIPR2-NEFH group is a distinct population of motor neurons in our samples, despite being largely defined by an enrichment of cytoskeletal genes

All six mixed clusters were spread throughout the ventral horn in laminae VII – VIII. MIXED-ADCYAP1 and MIXED-GCGR-TRIM54 showed no clear spatial organization, whereas MIXED-PTPRT-SALL3 neurons were found throughout the ventral horn and formed a band across dorsal lamina VII. MIXED-IL11-DCHS1 and MIXED-GABRA1-FXYD6 contained cells in laminae VII and VIII, plus some cells on the boundary of lamina X. The MIXED-FLT3-USP9Y cluster showed no obvious differences in spatial organization between males or females, noting that *USP9Y* transcripts were obviously not used to detect this population in female samples. This provides more evidence that this is not a male-specific population despite *USP9Y*, a Y chromosome-linked gene, being the most differentially expressed gene in the cluster.

All three of the inhibitory clusters found in this study (GABA-INSRR-NTRK1, GABA-SLC6A5-CSRNP1 and GABA-GCGR-DUOX2) had cells in laminae VII - XI. Consistent with rodent literature (*58*), the number of neurons in the glycinergic cluster appeared to be more numerous than those in GABAergic clusters in the ventral horn.

### Sex differences in human spinal cord microglia and astrocytes

There is a large body of evidence demonstrating sex differences in underlying mechanisms of chronic pain in preclinical animal models, and many of these differences are explained by glial cell-related mechanisms (*20, 21, 25, 59*). We therefore sought to assess whether there are sex differences in glial cell populations in the human spinal cord. In our snSeq analysis, astrocytes in the dorsal horn were subclustered into 15 distinct groups (Fig. 6A and B), while 13 subclusters were identified in the ventral horn (Supplemental Fig. 7). To further classify the astrocyte clusters by biological function, we grouped them into four categories: vascular, homeostatic, neurogenic, and metabolic gene-ontology enriched astrocytes (Fig. 6C). Homeostatic astrocytes were characterized by having enriched expression for genes involved in ion regulation and neurotransmitter clearance (*ATP1A2, SLC1A2, SLC6A11, AQP4*) (*60, 61*). Reactive astrocytes were defined by the enrichment of genes typically associated with injury or disease response (*MX1, CHI3L1, SERPINA3, IFIT1/2/3*)(*62, 63*). The metabolic cluster expressed canonical astrocytic markers but also showed strong enrichment for mitochondrial genes (*ATP6, CYB, MT-ND1–6, MT-CO1*), suggesting a heightened oxidative metabolic state previously noted in the literature (*64, 65*). These mitochondrial genes were not excluded from analysis, as their presence remained after several rounds of clean up, and likely reflect a genuine functional state of astrocytes rather than contamination. Neurogenic astrocytes demonstrated enrichment of genes associated with development, synaptic function, or a stem-like nature (*TNC, SPOCD1, DCLK2, GRIA2, NRCN1)* (*66, 67*). Finally, vascular-associated astrocytes showed enrichment of gene expression for transcripts linked to vascular interaction (*TMEM95, FOSB, PLVAP, ICAM4*).

**Figure 6.**
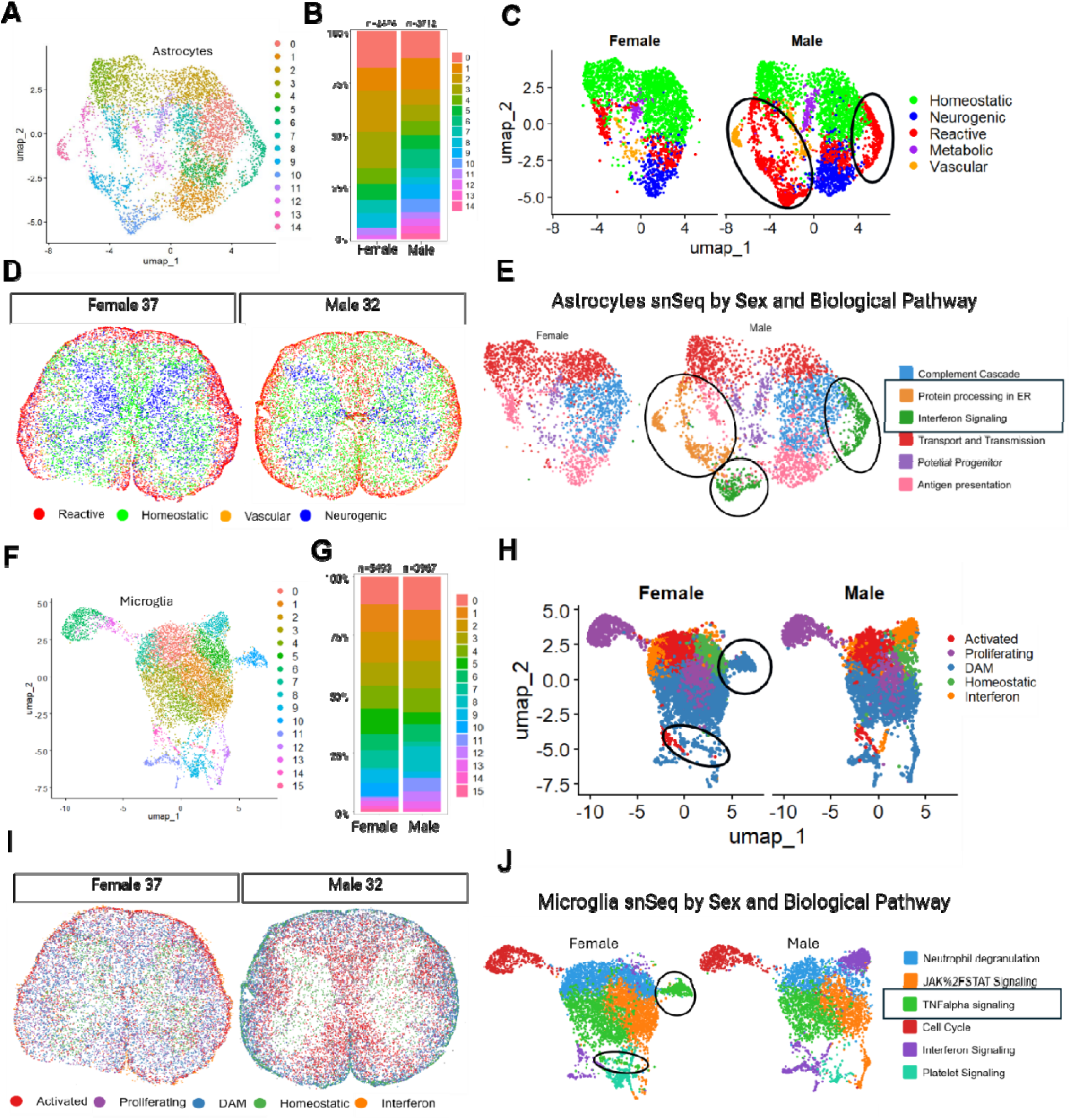
**Sex differences in human spinal cord microglia and astrocytes**. (**A)** UMAP of 14 single nuclei sequencing astrocyte clusters in the dorsal horn. (**B)** Stacked bar plot showing differences in astrocyte cluster proportions between males and females (**C**) UMAPs highlighting sex-specific expression patterns in astrocyte clusters; black circles indicate clusters, predominantly reactive, in males. (**D**) Spatial organization of astrocytes in representative male and female Xenium samples (**E)** UMAP of astrocytes annotated by biological pathway categories, showing male-enriched clusters involved in endoplasmic reticulum (ER) protein signalling and interferon signalling. (**F)** UMAP of 16 microglia clusters in the dorsal horn. (**G)** Stacked bar plot showing differences in microglia cluster proportions between males and females. (**H)** UMAPs showing sex-specific expression patterns in microglia clusters; black circles indicate clusters predominantly corresponding to disease-associated microglia (DAM) in females (**I)** Spatial organization of microglia in representative male and female Xenium samples. (**K**) UMAP of microglia annotated by biological pathway categories, showing female-enriched clusters involved in TNF_α_ signaling.

We examined potential sex differences and found that clusters 6, 9, 10, 13, and 14 were present only in males in the dorsal horn, with no sex-specific differences observed in the ventral horn (Fig. 6B, Supplemental Fig 7A). These clusters were predominantly classified as reactive, except for cluster 14, which was categorized as vascular. To investigate intercellular communication networks between astrocytes in both sexes, we analyzed the number and strength of interactions and observed stronger interactions between vascular and neurogenic astrocytes in females, whereas in males, the stronger interactions were between reactive and neurogenic astrocytes (Supplemental Fig. 7C and D). Next, using the subclustered Xenium astrocytes and their differentially expressed genes, we categorized them by function and mapped them back into a spatial context, as shown in the representative 32-year-old male and 37-year-old female samples (Fig. 6D). We were unable to identify a metabolic cluster due to the absence of genes representing this subtype on the Xenium panel. Similarly, we did not observe the same sexual dimorphism in the Xenium clusters as in the snSeq data, likely because the panel included a limited number of astrocyte-relevant genes. However, consistent with findings from snSeq, males appeared to have a greater number of reactive astrocytes that were diffusely distributed throughout the spinal cord, whereas in females, reactive astrocytes were more localized to the outer edges of the white matter (Fig. 6D).

Using GO term analysis of top-expressing genes within each astrocyte cluster, we identified six major biological pathways: complement cascade, protein processing in the ER, interferon signaling, transport and transmission, potential progenitor, and antigen presentation (Fig. 6E). Notably, the male-specific clusters showed predominant expression of genes involved in protein processing in the endoplasmic reticulum (e.g., *HSPA6, HSPA1A, HSPE1, DNAKB1*) and interferon signaling (e.g., *OAS1, OAS3, IFIT2, IFIT3, MX1, GBP3*). This finding contrasts with the DRG, where previous studies have shown a female-biased upregulation of type I interferon-related genes (*68*).

Turning to microglia, we identified 16 distinct clusters in the dorsal horn and 18 in the ventral horn (Fig. 6F). Similar to our approach with astrocytes, we categorized these microglial subclusters based on their putative functional states, resulting in five major groups: activated, proliferating, disease-associated microglia (DAM), homeostatic, and interferon-responsive.

Clusters exhibiting differential expression of genes involved in inflammation, complement activation, or immune modulation were classified as activated microglia (*C1QA, CD14, MRC1, LILRB5*) (69, 70). Clusters with elevated expression of cell cycle-related genes (*ZFP36L1, HIST2H2BE, CDK1, ANLN*) were designated as proliferating microglia (*71, 72*). Those expressing type I interferon response genes (*IFIT1, RSAD2, CXCL10, IFIT2*) were categorized as interferon responsive. Homeostatic microglia were identified by the expression of canonical markers including *CX3CR1, P2RY12, A2M, and MCF2L* (73). Finally, clusters expressing genes associated with neurodegenerative or dysfunctional states such as *CSF2RA, ITGAX, FTH1, MS4A7, GPNMB, LGMN, MARCO,* and *CD44* were classified as DAM-like microglia (*74, 75*). Two DAM clusters were exclusively observed in female samples (Fig. 6G-H). Spatial mapping revealed that in females, DAM cells were distributed throughout the white matter, whereas in males, they were more localized to the outer edges of the white matter. Further pathway analysis of the top-expressing genes in each cluster showed that the female-specific DAM clusters were enriched for TNFα signaling pathways. We categorized these microglial clusters based on relative gene expression and inferred function. Microglia are highly dynamic cells that shift their phenotype in response to environmental cues, and their functional states are often defined by surface marker expression, which we could not assess directly due to the use of single-nucleus rather than single-cell sequencing. Therefore, our classifications are based on transcriptional profiles and should be interpreted appropriately given the limited sensitivity of the single-nucleus approach in fully resolving microglial activation states.

### Spinal cord neuronal repertoire is adjusted to meet anatomical and evolutionary needs

Comprehensive human single-cell datasets provide an opportunity to explore how spinal cord neuronal repertoire changes through evolution and is adjusted to meet distinct functional needs along the rostro-caudal axis of the organ. To address this, we first generated a single-nucleus RNA-seq dataset from male and female mouse lumbar spinal cords (n=4 mice, male and female) yielding a dataset of 10,198 neurons. Furthermore, to assess cell-type level changes along the longitudinal axis of the human spinal cord we compared our human lumbar dataset to a recently published high quality cervical spinal cord atlas (n = 13,942 neurons(*9*)). Here, we focused on glutamatergic and GABAergic cells, as several recent studies have reported on cholinergic spinal cord neural diversity(*41, 76*).

We used canonical correlation analysis to align the three datasets in a joint embedding and identified orthologous neuron types (Fig. 7A). The increased cellular sampling across datasets allowed us to clearly distinguish 50 transcriptomic neuron classes where most directly corresponded to our previously identified human lumbar spinal cord neuron types or their subsets (Fig. 2,3). Importantly, 49 of the 50 neuron classes were clearly recognizable across species as well as across rostro-caudal segments of the human spinal cord (Fig. 7A-C) suggesting that the core neuron type architecture is strongly conserved throughout mammalian evolution and maintained along the longitudinal axis of the structure. However, we observed a high degree of variability in the relative abundance of individual neuron types when comparing spinal neuron types between homologous regions of the mouse and human spinal cord (stdev. of change from equal proportion = 18.56% with 17% of neuron classes showing three-fold enrichment as compared to the reference, Fig. 7B). This was in contrast to much lower changes in cell-type ratios across distinct regions of the human spinal cord (stdev of change from equal proportion = 11.67%, and 4% of neuron types displaying 3-fold enrichment, respectively, Fig. 7C) suggesting that altering the prevalence of individual transcriptomic neuron types is a common mechanism to meet evolutionary challenges impinging on these circuits.

**Figure 7:**
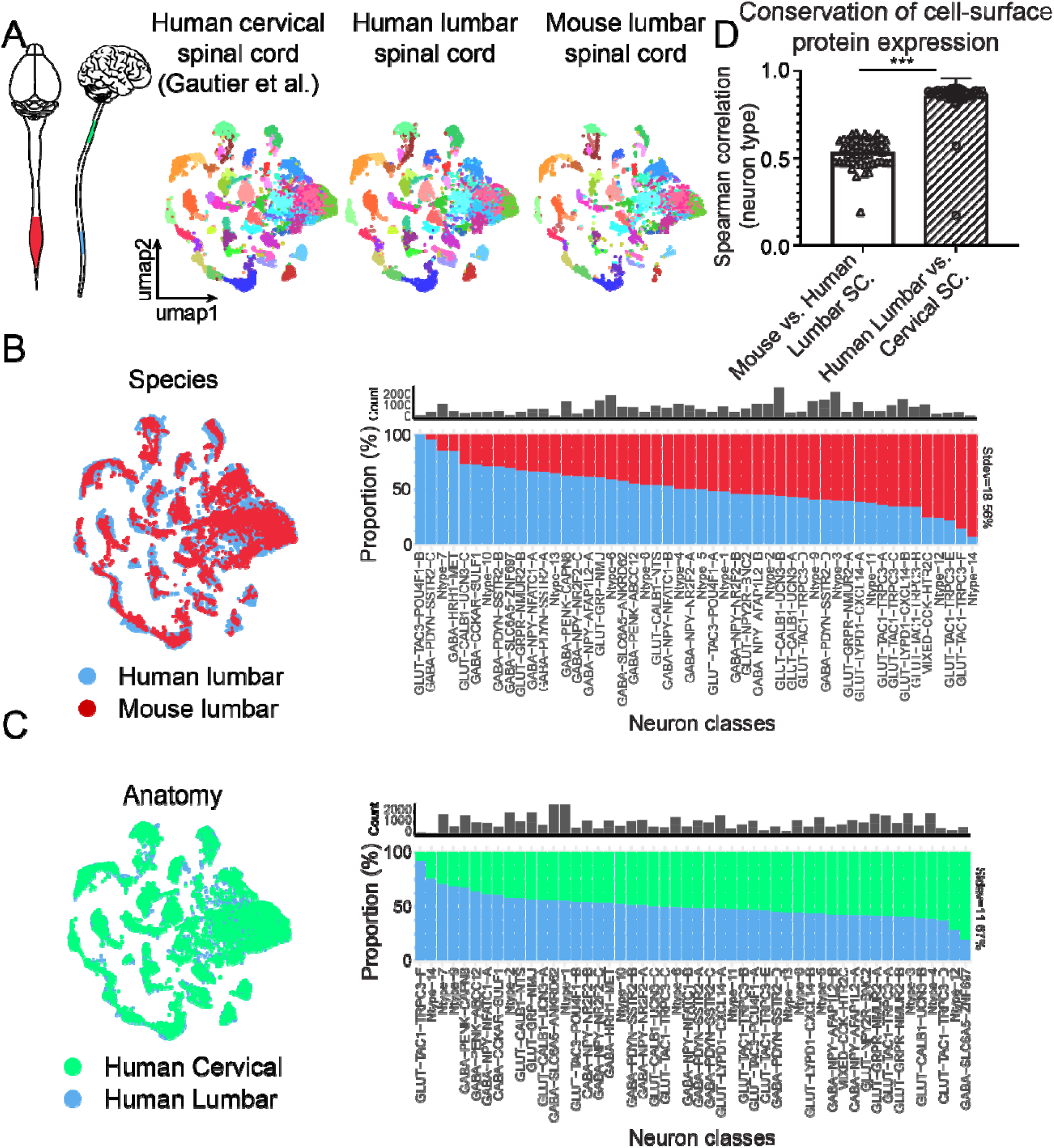
Anatomical and evolutionary variation in spinal cord neuronal repertoire. **(A)** UMAP embedding of aligned human cervical (n= 13,942 neurons), human lumbar (n=10 198 neurons) and mouse lumbar (n=12 696 neurons) spinal cord snRNA-seq data reveals orthologous neuron classes across anatomy and evolution. (**B)** Umap embedding of mouse lumbar (red) and human lumbar (blue) snRNA-seq datasets (left); Proportion of individual neuron types in an equally sampled mouse lumbar (n=10 198, red) and human lumbar (n=10 198, blue) snRNA-seq data (right), (**C)** Umap embedding of human cervical (from Gautier et al. 2023, green) and human lumbar (blue) snRNA-seq datasets (left); Proportion of individual neuron types in an equally sampled human cervical (n=12 696, green) and human lumbar (n=12 696, blue) snRNA-seq data (right), (**D)** Spearman correlation of the expression of top 315 most variable cell-surface genes between orthologous neuron types in mouse and human spinal cord or distinct human spinal cord subregions (two-tailed unpaired t-test, p<0.0001).

An alternative strategy for neurons to adapt to different species-specific functional requirements is to alter their molecular properties. These changes in individual cell-type molecular profiles can pose significant challenges in translating therapeutic approaches from model systems to human applications and are thus important to understand (*32, 33*). To this end, we evaluated the correlation of cell-surface proteins encoding genes across individual classes of spinal cord neurons contrasting mouse to human and human cervical to lumbar spinal cord differences. Predictably, we observe a dramatically higher correlation between most cell-surface proteins in corresponding cell types between different human spinal cord domains as compared to corresponding spinal cord neuron classes between human and mouse (mean Spearman correlation = 0.76 +/- 0.10 vs. 0.53 +/- 0.09, respectively, data shown as mean +/- standard deviation, p < 0.0001, Fig. 7D, Supplemental Table 3). Collectively, these data argue, that the neuron type repertoire is remarkably stable across the longitudinal axis of the spinal cord as well as throughout mammalian evolution. However, the precise ratio as well as molecular features of individual neuron types are different between humans and mice.

### Electrophysiological characterization of human lamina II dorsal horn neurons

To better understand the physiology of human dorsal horn neurons, we developed a method to obtain acute parasagittal slices from post-mortem human spinal cord (Fig. 8A) and performed whole-cell recordings from neurons in lamina II (LII) of the dorsal horn (Fig. 8B). Human LII neurons exhibited qualitatively similar firing patterns compared to rodents in response to depolarizing currents (Fig 8C), which were predominated by “tonic” (5/16) and “transient” (9/16) firing phenotypes (Fig. 8D). Tonic firing neurons exhibited a high firing frequency in response to supratheshold stimuli, while transient neurons exhibited action potential accommodation and reduced firing frequencies (Fig. 8E). We compared a number of active and passive membrane properties, including resting membrane potentials, input resistance, rheobase and AP voltage thresholds, which were not different between these two types (Fig. 8F, Supplemental Fig. 11A-G). However, one third of transient neurons (3/9) exhibited a shoulder on the AP falling phase, represented by a hump in the first derivative plots (Supplemental Fig. 11I-K), which we and others have observed previously in large proportion of human presumptive C-fiber DRG neurons (*77–79*) but has not been reported previously in rodent dorsal horn. Tonic firing neurons did not exhibit this shoulder (Supplemental Fig. 11) and had a higher proportion of spontaneous activity at rest (Fig. 8G, H). Using biocytin cell fills, we reconstructed morphology from a subset of these recordings and observed that tonic firing neurons had larger dendritic arbors mostly confined to LII and extending in the rostral-caudal direction (Fig 8I), reminiscent of islet cell morphology from rodents (*80, 81*). In contrast, the dendritic branching of transient firing neurons were much smaller and exhibited minimal rostral caudal elongation (Fig 8I), reminiscent of radial type neurons observed in rodents.

**Figure 8.**
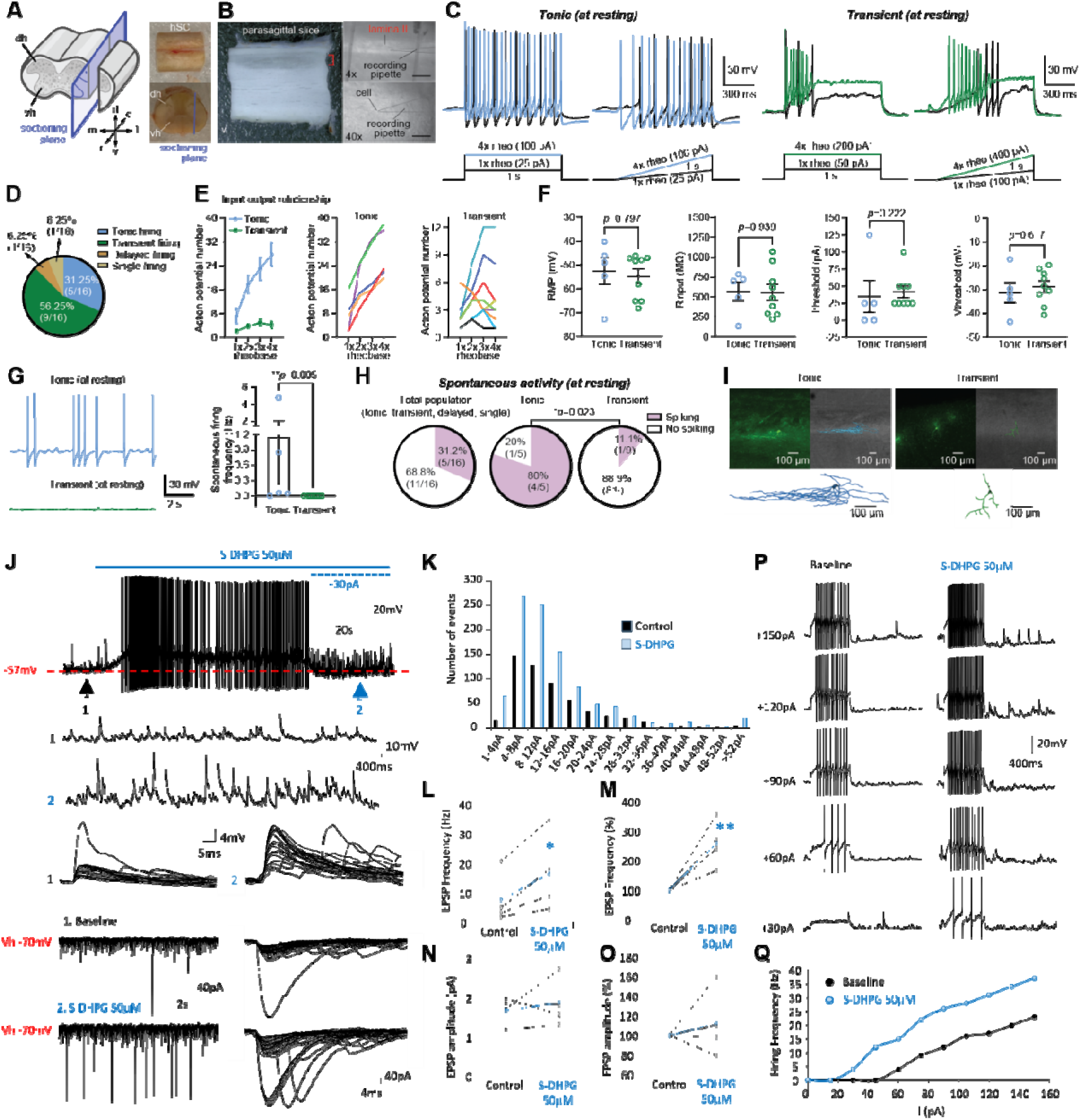
**Functional properties of human dorsal horn neurons and sensitization by mGlu1/5 agonists**. (**A**) Schematic illustrating the sectioning orientation of the hSC to obtain the sagittal slices (left) and fresh hSC tissue samples (right). (**B**) Targeted whole-cell patch-clamp recordings of lamina II hSC slices with representative differential interference contrast (DIC) images at 4x and 40x magnification. Scale bar: 445µm and 45µm respectively. (**C**) Representative voltage responses of tonic (left) and transient (right) firing hSC lamina II neurons recorded at resting membrane potential. Neurons were stimulated with depolarizing current steps and ramps at 1x and 4x rheobase (rheo). (**D**) Pie chart summarizing the distribution of firing patterns among recorded hSC lamina II neurons, including tonic firing, transient firing, delayed firing, and single firing (n=16 cells/5 donors). (**E**) Input-output (I-O) relationship of hSC lamina II neurons. Left: Average action potential number elicited by increasing rheobase current for tonic (n=5) and initial (n=9) firing neurons. Middle and Right: Individual I-O curves for all recorded tonic and initial firing neurons. (**F**) Comparison of intrinsic properties between tonic and initial firing neurons, including resting membrane potential (RMP, unpaired two-tailed Mann-Whitney test), input resistance (Rinput, two tailed unpaired t-test), rheobase (Ithreshold, unpaired two-tailed Mann-Whitney test), and voltage threshold (vthreshold, two tailed unpaired t-test) (**G**) Spontaneous firing activity at resting membrane potential. Left, representative traces showing spontaneous action potentials in tonic but not initial firing neurons. Right, quantification of spontaneous firing frequency in tonic and initial firing neurons (unpaired two-tailed Mann-Whitney test). (**H**) Pie charts summarizing the proportion of spontaneously spiking versus non-spiking neurons in the total population and within tonic and initial subgroups (tonic vs. initial, Fisher’s exact test). (**I**) Micrographs of neuronal morphology from biocytin cell fills (top) and reconstructions (bottom) for tonic and transient firing cell types. (**J**) Samples of a continuous recording showing S-DHPG induced membrane potential depolarization, action potential discharge and an increase in spontaneous excitatory post-synaptic potentials (EPSPs) in this neuron, the latter revealed by injection of negative current (−30pA) at the peak of the response. (**L**) Amplitude histograms illustrating increased frequency of events on spontaneous EPSCs in S-DHPG treated cells. (**L** to **O**) Plots showing the effects of S-DHPG on the frequency of spontaneous EPSPs in all neurones (n=4). S-DHPG induced a significant increase in the frequency of EPSPs when data were expressed as absolute values (p<0.05) and with each cell normalised to its own baseline spontaneous EPSP frequency (p<0.01; paired Student’s t-test). There was no significant effect on spontaneous EPSP amplitude when expressed as absolute values or with data normalised to their own baseline EPSP amplitude. Note the blue plots represent the mean ± sem of the population (n=4). (**P**) Recording and (**Q**) plot of membrane responses to linearly increasing depolarising rectangular-wave current pulses in the absence (Left column) and presence (right column) of S-DHPG demonstrating an increase in intrinsic electrical excitability of dorsal horn neurons.

### Group I mGluR agonist-induced excitation and sensitization

We found that GRM5, which encodes the group I metabotropic glutamate receptor mGlu5, was among the most highly expressed excitatory GPCR among dorsal horn neurons (Supplemental Fig 2F). Therefore, we sought to assess plasticity induced by an agonist of this receptor on human dorsal horn neurons. The effects of the group I mGluR receptor agonist S-DHPG were investigated on 9 lamina II neurons in spinal cord slices prepared from 3 donors. S-DHPG (30-50 μM) induced membrane potential depolarization in 8 neurons and was without any notable effect in 1 neuron consistent with lack of GRM5 expression in some dorsal horn neurons. S-DHPG-induced significant membrane potential depolarization which was sufficient to reach threshold for action potential firing in 5 neurons (Fig. 8J). S-DHPG-induced membrane potential depolarization was also associated with concomitant increases in spontaneous excitatory post-synaptic potentials in 5/8 neurons that responded (Fig. 8J). In the presence of S-DHPG, spontaneous EPSPs showed a significant increase in frequency and a non-significant trend toward increased amplitude (Fig 8K to O). Given the lack of expression of group I mGluRs in hDRG (*82*), this suggests a presynaptic action on excitatory interneurons of the dorsal horn. In 3 neurons, S-DHPG-induced depolarization was associated with an increase in spontaneous inhibitory post-synaptic potentials (IPSPs) or currents (IPSCs), the latter observed in voltage clamp mode at a holding potential of −40Mv. Action potential firing frequency, evoked in response to depolarizing rectangular-wave current pulses, was also enhanced in the presence of S-DHPG compared to baseline (Fig. 8P and Q).

## Discussion

In this study, we have deeply sequenced the dorsal and ventral regions of the human lumbar spinal cord using spatial technology with a single-molecule resolution to determine precise neuronal organization and reveal sex differences in glial cell clusters. We have also compared this human lumbar snSeq data to a previously published human cervical spinal cord snSeq dataset and a generated snSeq dataset of mouse lumbar spinal cord, revealing conservation of cell types but evolutionary changes in cell surface proteins. Electrophysiological recording support conservation of basic physiological properties with what is known in mice. Finally, consistent with widespread expression of the GRM5 gene across dorsal horn neuronal cell types, we observed plasticity in dorsal horn circuits upon stimulation with S-DHPG.

### A comprehensive map of human spinal cord neurons

Several previous studies have sequenced entire human spinal cord sections and have therefore required a combination of gene expression and deconvolution of low-resolution spatial sequencing data to separate dorsal and ventral horn populations (*8, 10, 12*). In this study, we sequenced the two regions separately and used single-molecule spatial technology to confidently place neurons in space and gain further clarity on smaller ventral horn populations that have been previously understudied or the topic of controversy in the field (*7–9*). In total, we identified 20 dorsal horn and 14 ventral horn populations. We found that neurons in the dorsal horn showed a highly specific laminar distribution, with many recapitulating rodent orthologs implicated in pain processing (*83*). For example, the GLUT-GRPR-NMUR2 population was restricted to lamina II and displayed high expression of various peptide receptors, likely because these neurons are densely innervated by peptidergic primary afferents, potentially mirroring the *Grpr*-expressing vertical cells described in rodent (*84*). In the rodent, vertical cells receive inhibitory input from parvalbumin-containing inhibitory interneurons, which gate low threshold mechanoreceptors via presynaptic inhibition (*85*). We find that *PVALB* is expressed in one of the dorsal glycinergic populations, GABA-SLC6A5-ANKRD62, suggesting a similar circuit may underlie mechanical pain sensitivity in humans. In our view, the current findings set a foundation of cellular understanding of the human dorsal horn that can be used across future studies to predict and unravel the circuitry underlying gating of nociceptive and mechanical sensory input in the human spinal cord.

A single human and several rodent transcriptomic studies have dissected and sequenced the ventral horn separately from the dorsal, and all of these have focused on CHAT-expressing motor neurons rather than ventral horn interneuron clusters (*12, 14, 76, 86–88*). Isolating ventral horn cells from the entire spinal cord via gene expression or spatial sequencing with a low-resolution has been widely used in rodent and some human studies (*8, 10, 12, 41, 89*), however this may not accurately capture all ventral horn neuronal populations. We have identified 11 interneuron populations in the ventral horn; however, it is hard to place many of these into a broader, cross-species context due to a lack of knowledge regarding ventral horn interneurons in comparison to the dorsal horn. Our data reveal separate alpha and gamma motor neuron groups, together with a third motor neuron cluster similar to that observed by Yadav et al., (2024) and marked by high expression of cytoskeletal genes. Furthermore, we identified large motor neurons with marker genes of this population in spinal cord sections processed with Xenium spatial transcriptomics. Future studies can build on these findings using disease-related samples, to further dissect ventral horn circuity.

### Implications of sex differences in human spinal cord microglia

Our snSeq and Xenium spatial transcriptomic analysis revealed clear sex differences in spinal glial transcriptional states, including astrocytes, microglia, and oligodendrocytes, suggesting sexual dimorphic cellular strategies in modulating neuro-immune responses (Fig. 6, Supplemental Fig. 9). These findings align with previous reports indicating sex-dependent variation in neuroinflammatory responses and glial activation states in the context of neurological diseases such as ALS and chronic pain(*19, 90, 91*). Specifically, we identified two microglial subpopulations present exclusively in females, enriched for signatures of disease-associated microglia (DAM) and characterized by upregulation of TNFα signaling pathway genes. Previous rodent studies have suggested a male-dominant role for certain types of microglial signaling in the development and maintenance of neuropathic pain in the dorsal horn(*92–94*), but other studies have shown that microglia can promote chronic pain in female mice, albeit with far less understanding of the underlying mechanisms(*26, 95*). The identification of female-exclusive DAM-like microglia population enriched for TNFα-mediated signaling suggests that this population of CNS immune cells might contribute to synaptic plasticity in the dorsal horn(*96–98*) that is associated with central sensitization(*16*). Future studies on dorsal horn samples from individuals who died with a history of chronic pain can shed light into how these populations of microglia might change in sex-specific ways in humans with chronic pain.

We identified two oligodendrocyte subpopulations in the ventral horn and five astrocyte subpopulations in the dorsal horn that were male-specific. In the male-exclusive oligodendrocyte clusters, we found several top-expressing genes with links to ALS, including *SQSTM1*, *HSPB1*, and members of the HSP family chaperones. These genes are involved in proteostasis, stress response, and protein aggregate clearance (Supplemental Fig. 9)(*99–101*). Elevated baseline expression of these pathways may reflect a sex-specific compensatory adaptation to increased metabolic demands required to support neuronal function and survival. This intrinsic difference could contribute to the observed male vulnerability in ALS, potentially rendering male oligodendrocytes more susceptible to pathological overload as the disease progresses(*102*).

Likewise, the male-specific astrocyte clusters exhibited a reactive transcriptional profile with upregulation of interferon response pathways, which have been implicated in chronic pain both in the CNS (*103, 104*) and in the periphery (*68, 105*). Mouse studies suggest that type I interferon signaling in the dorsal horn suppresses pain signaling (*104*), with more powerful effects in male mice (*103*), so this population of astrocytes may function to decrease nociceptive processing in the dorsal horn specifically in men. More broadly, spinal astrocytes have gained attention due to their involvement in the development and maintenance of pain, particularly the reactive astrocytes which secrete inflammatory cytokines that can interact with microglia and alter neuronal excitability (*106*). Therefore, these male-specific reactive astrocyte clusters may reflect an alternative axis of glial reactivity in males, potentially engaging neuroimmune signaling cascades distinct from those observed in females.

Notably, one of the male-specific astrocyte clusters exhibited a vascular-related transcriptomic profile (Fig 6). Astrocytes form gap junctions that interface with the vasculature, and inhibition of these gap junctions have been shown to reduce pain hypersensitivity in mouse neuropathic pain models (*107, 108*). Given the critical role of astrocytes in maintaining neurovascular unit integrity, this observation led us to investigate potential sex differences in spinal cord ependymal cells and endothelial cells (Supplemental Fig. 10)(*109, 110*). While we did not detect any sex differences in ependymal or endothelial populations, we observed an age-associated increase in endothelial cell density and vascular penetration into the spinal gray matter, particularly of angiogenic (Tip-like) endothelial cells (Supplemental Fig. 10). This contrasts with the widely reported vascular rarefaction seen with aging (*111, 112*). Our spatial transcriptomic data suggests that the spinal cord follows a distinct aging trajectory that warrants future investigation. These findings have important implications for age-related spinal vulnerability, neuroimmune regulation, and drug accessibility across the lifespan. More broadly, our results highlight sex-specific glial specialization in the spinal cord, providing a cellular framework for divergent neuroimmune mechanisms underlying pain processing.

### Differences and similarities in spinal cord neuronal populations across species

Despite 85 million years of evolutionary divergence between mice and humans (*113*) we and others (*8, 12*) have found extensive homology between their spinal cord neuronal repertoires. We analysed how human spinal cord cellular diversity is adjusted to meet distinct functional requirements along the rostrocaudal axis. Somewhat surprisingly, we failed to find any rostro-caudally specific GABAergic or glutamatergic neuron classes, which is in notable contrast to the spinal cord cholinergic system with many neuron types with highly restricted rostro-caudal locations such as different visceral motor-neuron types (*41, 76*). Upcoming studies may however uncover extensive rostrocaudally unique neuron types in other spinal cord regions including thoracic and sacral spinal cord, which were not analysed in the current work.

We also characterized the mechanisms by which orthologous neuron types differ across mice and humans. Despite finding little evidence for species-specific neuron types, we observed extensive changes in the relative abundance of orthologous neuron classes in corresponding anatomical spinal cord regions. Furthermore, we also characterized moderate correlation in differentially expressed cell surface genes in homologous neuron types in mice and humans. This likely partially explains the high failure rate in translation of pre-clinically successful therapeutic applications to human therapies (*114*) and suggests that for some conserved cellular targets different molecular pathways may need to be therapeutically tapped. An example of a well-conserved neuronal class and neuropeptide to transmitter receptor pair is the gastrin releasing peptide (GRP) and GRP receptor (GRPR) system that has been extensively studied in the spinal cord in the context of chemically induced itch. We found that *GRP* and *GRPR* marked specific subtypes of neurons in the human dorsal horn in equivalent spatial locations in lamina II to what has been described in mice (*115*). This supports the importance of these subsets of neurons in chemical itch in mice and humans and highlights their value as a therapeutic target for chronic itch diseases (*116, 117*). Likewise, we found that the NPY1R was expressed in populations of neurons that have previously been linked to neuropathic pain relief in mice by either chemogenetic inhibition or agonism of this receptor. However, we also noted a broader expression of the receptor in excitatory interneurons in humans than has been appreciated in previous studies in mice (*118, 119*). Our extensive characterization of receptor expression in populations of spinal cord neurons should enable translational decisions about targets that are under consideration for pain treatment, or other spinal cord diseases.

Another example of cross-species conservation is found in our physiological recordings from the spinal cord. Here we found similar firing patterns among dorsal horn neurons to what has previously been observed in rodents(*83*), although we did note a species difference in action potential shape that likely reflects differences in gene expression for cell surface ion channels, consistent with our cross-species transcriptomic analysis. We also found group 1 mGluR plasticity that is consistent with previous observations in rodents. In the mouse dorsal horn, activation of group 1 mGluRs causes central nociceptive sensitization (*120*) and mGluR5 activation produces increased excitability of these neurons via modulation of Kv4.2 (*121*). These previous studies align with our findings in human dorsal horn recordings and validate mGluR5 antagonism as target for manipulation of pain in humans.

### Conclusions and future directions

Our work provides unprecedented insight into the cellular composition, in spatial context, of the human dorsal and ventral horn. Our findings provide a grounding for investigation of human spinal neuronal circuits based on neuronal subtype, homology to mouse where extensive circuit mapping has already been done, anatomical location within the laminae of the spinal cord, and physiological properties. We also reveal sex differences in glial subtypes that are likely relevant for disease states in humans. An important future direction will involve building on this foundation to identify dorsal horn neurons that make up the spinothalamic tract in humans, the only route for nociceptive information to reach the brain, where pain perception occurs. Our findings suggest that the GLUT-TAC3-POUF4F1 population may contain this critical population of neurons. Another important area will be to use methods established herein to interrogate the human spinal cord in disease states such as neuropathic pain, or motor diseases that affect lower motor neurons, like ALS.

## Methods

### Human Spinal Cord Tissue retrieval

All human tissue procurement procedures were approved by the Institutional Review Board at the University of Texas at Dallas under protocol Legacy-MR-15-237 and collected in collaboration with the Southwest Transplant Alliance. The Southwest Transplant Alliance obtained informed consent for research tissue donation from first-person consent (driver’s license or legally binding document) or from the donor’s legal next of kin. Policies for donor screening and consent are those established by the United Network for Organ Sharing (UNOS). OPOs follow the standards and procedures established by the US Centers for Disease Control (CDC) and are inspected biannually by the Department of Health and Human Services (DHHS). The distribution of donor medical information is in compliance with HIPAA regulations to protect donor privacy.Spinal cords from the thoracic to the sacral level were surgically dissected from human organ donors within 1.5 - 3.5 hours of cross clamp and immediately frozen using crushed dry ice as previously described(*122*). Blocks were stored at −80°C until use (see Table 1 for donor demographic information).

### Generation of human spinal cord nuclear suspensions

The detailed protocol has been previously published(*123*). In summary, the key steps required dividing the dorsal horn and the ventral horn and extensive cleaning off the outer pia layer of the fresh frozen lumbar spinal cords. The tissues were homogenized in lysis buffer using 5 strokes with the loose pestle followed by 15 strokes with the tight pestle. Homogenate was filtered using a 70-micron cell strainer on a 50 mL Falcon tube and pre-wet with Nuclear Buffer. Nuclei were spun for 5 min at 500g at 4°C in a spin-out rotor. Supernatant is removed and pellets were resuspended in nuclear buffer and gently layered onto 12 mL of Sucrose Cushion in a 50 mL Oakridge tube and centrifuged at 3200g at 4°C for 20 minutes in a spin-out rotor. After centrifugation supernatant was removed by decanting in one smooth motion and drying out the neck of the Oakridge tube with a Kimwipe. At this point the nuclei were either resuspended using 1mL of fixation buffer from the 10X fixation of cells & nuclei for chromium fixed RNA profiling (CG000478) and transferred to a 15mL Falcon tube and incubate for 18hr at 4°C. The nuclei were centrifuged at 500g for 5min at 4°C and resuspend in Nuclear Buffer if FACS sorted or directly incubated in quenching buffer and continue with sequencing. For FACS sorting, a previously conjugated anti-NeuN-Alexa-647 Ab and 7-AAD were added to the nuclear suspension and incubated in the dark for 15 minutes. Nuclei washed and spun at 500g for 5 min at 4°C in a spin-out rotor twice. Remove supernatant, resuspend the nuclei and gently triturate with 150-micron glass Pasteur pipette to de-clump any nuclei. The nuclei were sorted at low pressures (4 or 5 PSI on BD sorters) and selected for 7-AAD and Alexa 647 double-positive nuclei after compensation with single-stains of dyes.

### Single-nucleus RNA sequencing for human samples

Single-nucleus RNA sequencing was carried out using the Chromium Next GEM Single Cell Fixed RNA Sample Preparation kits from 10XGenomics and used according to the manufacturer’s instructions. The library was created using 10x Fixed RNA Gene Expression using a NextSeq1k2k instrument platform (v3.8.4) at the Genome core at the University of Texas at Dallas. Libraries were sequenced to a minimum depth of 20,000 reads per nucleus using an Illumina HiSeq 3000 (PE 26–8 – 98 bp). Raw sequencing reads were demultiplexed, aligned, and a count matrix was generated using CellRanger (v6.1.2). For alignment, introns and exons were included in the human reference genome (GRCh38).

#### Generation of mouse spinal cord nuclear suspensions

8-week old male and female mice were anesthetized with isoflurane and decapitated. Vertebral column was rapidly dissected out and transferred to ice-cold carbogenated (95% O_2_ and 5% CO_2_) NMDG-HEPES-ACSF (93 mM NMDG, 2.5 mM KCl, 1.2 mM NaH_2_PO_4_, 30 mM NaHCO_3_, 20 mM HEPES, 25 mM glucose, 10 mM MgSO_4_, 1 mM CaCl_2_, 1 mM kynurenic-acid Na salt, 5 mM Na-ascorbate, 2 mM thiourea and 3 mM Na-pyruvate, pH adjusted to 7.4, osmolarity ranging 300–310 mOsm). L1-L5 segments of the spinal cord were microdissected out and cut into small 1 mm^2^ size pieces. Microdissected spinal cord pieces were transferred to a Kimble Dounce tissue-grinder containing 750 microL of Lysis Buffer (0.1% Triton-X100, 320 mM sucrose, 10 mM Tris-HCl – pH=7.4, 3mM MgCl_2_, 10 mM NaCl, 0.5% RNAse free BSA, 1% Kollidon VA64, 1mM dithiotreitol, 0.1 U/microL Roche Protector RNAse inhibitor in DNAse/RNAse free ddH_2_O). Tissue was homogenized with 5 strokes with loose pestle and 15 strokes with tight pestle and resulting nuclear suspension was strained with a 40 micron cell strainer and washed with 1.25 mL of Nuclear Buffer (320 mM sucrose, 10 mM Tris-HCl – pH=7.4, 3mM MgCl_2_, 10 mM NaCl, 0.5% RNAse free BSA, 1% Kollidon VA64, 1mM dithiotreitol, 0.1 U/microL Roche Protector RNAse inhibitor in DNAse/RNAse free ddH_2_O). Cell suspension is layered onto 12 mL of 1M sucrose cushion (1M sucrose, 10 mM pH=7.4, Tris-HCl (pH=7.4), 3mM MgCl_2,_ 10 mM NaCl, 0.5% RNAse free BSA, 1% Kollidon VA64, 1mM dithiotreitol, 0.1 U/microL Roche Protector RNAse inhibitor in DNAse/RNAse free ddH_2_O) in a 5 mL Oakridge tube. The latter are centrifuged at 500g for 5 min at 4C in a spin-out-rotor. The sucrose cushion is decanted and pure nuclei are resuspended in 300 µL of nuclear buffer containing 10 µg/mL 7-AAD and 1:100 custom conjugated anti-NeuN-Alexa647 antibody (Biolegend #608452). The nuclei are incubated for 15 min at 4C. Thereafter, nuclei are spun down at 500g for 5 minutes and washed twice with 3 mL of nucleus buffer and resuspended in 1 mL of nucleus buffer. The resulting nucleus suspension is FACS sorted to enrich for NeuN/7-AAD double positive cells (neurons). The neuronal nuclei are concentrated with CUTANA ConA beads per manufacturer’s protocol and resuspended in 1x Nucleus Buffer from 10x Genomics with resulting nuclear suspension loaded to the 10x Genomics microfluidic chip and processed per manufacturer’s protocol.

### Single-nucleus RNA sequencing for mouse samples

Single-nucleus suspensions were processed on a 10X Chromium Xi instrument using the Chromium Next GEM Single Cell Multiome Reagent Bundle (#100028) following the manufacturers protocol. The resulting sequencing libraries were sequenced on a Novaseq X platform (PE 150 bp) at Admera Health. Libraries were sequenced to a minimum depth of 50,000 reads per nucleus. Raw sequencing reads were demultiplexed, aligned, and a count matrix was generated using CellRanger Arc (v2.0.2) using the Mouse mm10 (GENCODE vM23/Ensembl98) reference transcriptome with –include introns parameter specified.

### Computational analysis

We used Rstudio (v4.3.2) and Seurat (v5.3.0) for data analysis. We created all 11 objects with a minimum of 3 cells and minimum 200 features. We then normalized the data and used the vst as a selection method with 2000 features. FindIntegrationAnchors function was used with dims 1:10. We the integrated the data to create one larger object. This was then subset for greater than 500 features and less than 5 percent mitochondrial genes. The data was then scaled using the ScaleData function followed by principal component analysis (PCA). We used the ElbowPlot function to determine the final parameters and ran RunUMAP and then FindNeighbors with pca reduction and dims 1:15. Next we used FindClusters with a resolution of 1. Highly similar clusters without clearly distinguishable markers were merged to produce the final 30 clusters for the dorsal horn and 27 clusters for the ventral horn.

### Cell type annotation and reclustering

Dorsal and ventral horn clusters were analyzed separately. Following quality control, remaining cells were grouped into 30 clusters for the dorsal horn (from 97,045 cells) and 27 clusters for the ventral horn (from 81,996 cells). Differentially expressed genes (adjusted p<0.05) of each cluster in comparison to other clusters were manually searched for well-defined cell type markers and assigned a cell type. Clusters of each cell type were combined to leave 8 populations: oligodendrocytes and their precursors, neurons, astrocytes, microglia, endothelial cells, lymphocytes, fibroblasts and ependymal cells. Each of these groups was then separately re-clustered to reveal subpopulations of each cell type. We performed a manual cleanup of doublets based on well-established markers for each cell type (for neurons: *MOG, RBFOX3* Oligodendrocytes: *MOG, MYRF* Astrocytes *AQP4, ALDH1L1* Endothelial: *VWF, ABCB2* Microglia: *CSF1R, CLEC7 Fibroblasts: C7, COL3A1,* Lymphocytes: IL2RG, CXCR4, Ependymal: FOXJ1, CFAP157) and re-clustered several times until we determined the cleanest subclusters.

### Cell Chat and Interactome

To assess intercellular communication between cell types, CellChat package (version 1.6.1) which contains a comprehensive interaction database of receptor-ligand interaction was used. It uses probability based on the law of mass action to estimate the likelihood of communication between cell types. For statistical analysis, it uses permutation testing, to distinguish biological relevant interactions. We utilized this package to assess changes in interactions between male and female astrocytes.

### Neuronal subclustering and annotation

In the dorsal horn, 20 subpopulations were grouped from 11,457 neurons and 14 neuronal subpopulations were found from 2325 neurons in the ventral horn. Differentially expressed genes were again searched to annotate each population. A three tiered naming system was chosen to provide the greatest amount of information: the first name refers to the main fast neurotransmitter of the cluster such that GLUT clusters contain glutamatergic genes, GABA clusters are inhibitory and contain GABA and / or glycinergic genes, CHOL clusters contain cells with cholinergic genes and MIXED clusters have both excitatory and inhibitory nuclei. The second tier of the name refers to a differentially expressed gene that may be of interest to the field and could relate to signaling such as those encoding peptide precursor proteins. The final tier refers to the top differentially expressed gene of each cluster. In a single ventral horn cluster, only one gene was differentially expressed when compared to other ventral horn neuron populations and in this case the annotation is a two-tiered name only (MIXED-ADCYAP1).

To assess sex differences in neuronal populations, we looked at the proportion of neurons assigned to each cluster in males and females. To account for more neurons coming from female samples than male samples, we normalized for the total number of neurons per donor. These proportions were compared for each cluster using a non-parametric Wilcoxon rank test in R.

### Non-neuronal clustering

To investigate transcriptional heterogeneity among astrocytes, microglia, oligodendrocytes, endothelial cell and ependymal cells, we performed sub-clustering in Seurat (v5.3.0) using RStudio (v4.3.2). Each of the cell subtypes from the dorsal and ventral horns were subset from the integrated dataset based on prior cell-type annotations. Data were scaled and subjected to principal component analysis (PCA) using variable features. The ElbowPlot was used to determine the number of principal components to retain, and the top 15 PCs were selected for downstream analysis. Dimensionality reduction was performed using UMAP, and clustering was carried out with the Louvain algorithm (FindClusters, resolution = 1). To identify differentially expressed genes defining each cluster, we used the non-parametric Wilcoxon rank sum test to compare gene expression in each cluster against all other clusters using the FindAllMarkers function. Marker gene expression and gene ontology (GO) enrichment analyses were used to annotate clusters by biological function. For the functional categorization, we did a literature search for the most relevant genes across each functional state and cited in the text. Differential expression was calculated between male and female samples to identify sex-specific non-neuronal sub-populations. Mitochondrial genes were retained in the analysis of the astrocytes to capture functionally relevant metabolic states. Clustering and visualization steps were repeated independently for dorsal and ventral horn regions. The final Seurat objects were saved as .rds files for downstream analysis and Loupe file generation

### Xenium processing and analysis

Frozen blocks of lumbar spinal cord (n = 8, 4 males & 4 females; see Table 1 for donor demographics) were embedded in OCT by applying a small layer at a time and allowing it to freeze on dry ice before proceeding with the next layer until fully covered, to reduce thawing. These blocks were cut into 10µm transverse sections using a Leica cryostat and mounted onto Xenium Slides (cat # PN-1000460), as per 10X Xenium In Situ for Fresh Frozen Tissues – Tissue Preparation Guide guidelines (https://www.10xgenomics.com/support/in-situ-gene-expression/documentation/steps/tissue-prep-fresh-frozen/xenium-in-situ-spatial-profiling-for-fresh-frozen-%E2%80%93-tissue-preparation-guide). Slides were immediately placed in a −80°C freezer overnight and used the following day.

Slides were then processed following the Xenium In Situ Gene Expression with Cell Segmentation Staining User Guide (https://www.10xgenomics.com/support/in-situ-gene-expression/documentation/steps/assay/xenium-in-situ-gene-expression-with-morphology-based-cell-segmentation-staining-user-guide). All slides were incubated with a 480-custom gene Xenium spatial transcriptomics panel containing cell type and neuronal subpopulation marker genes and 10X Cell Segmentation Staining Reagents (cat # PN-1000661). The custom gene panel contained marker genes for each cell type and neuronal subcluster, together with receptors and ion channels of interest (see Supplemental Table 2 for a list of genes included). Some marker genes were not included on the panel due to high expression levels that would lead to optical crowding as per 10X recommendations, and the amount of amplification of each of the remaining genes on the panel was considered carefully in coordination with the single-nucleus sequencing data described above, to ensure high expressing genes were amplified fewer times than lower expressing genes.

### Neuronal spatial organisation in the human spinal cord

To map the neuronal clusters in space, we sought to find a combination of approximately 3 differentially expressed genes that together capture the largest proportion of each cluster, whilst capturing the fewest cells in other clusters. Some clusters required more genes to increase the specificity, whilst others required fewer than 3. The ventral MIXED-ADCYAP1 cluster only had one differentially expressed gene, *ADCYAP1*, to identify the cluster. As the ventral MIXED-FLT3-USP9Y cluster had a Y chromosome-linked gene as the most differentially expressed gene, this gene was only included to identify the cluster in males. In females the remaining marker genes of the cluster were used to identify this population (see Supplementary Table 2 or Supplementary Fig.3 to see marker genes for each population).

Following Xenium processing, the composite morphology images (DAPI + multi-channel immunofluorescence staining included in the 10X Cell Segmentation kit as follows: immunolabeling for ATP1A1, CD45, E-cadherin for cell boundaries in a single channel, 18S ribsosomal RNA for internal RNA in a second channel and alphaSMA and vimentin for internal protein in a third channel) were overlaid with canonical neuronal markers RBFOX3 and GRIN1 and glial/stromal markers AQP4, MOG, CSF1R and ABCG2. In QuPath v0.4.4, individual cells that expressed neuronal transcripts with no non-neuronal transcripts were interactively outlined with the polygon tool to generate high-fidelity boundaries. The curated objects were exported as GeoJSON, producing a single segmentation.geojson that contains one FeatureCollection. The ROIs were then reintegrated back into xenium data by running the xenium bundle through import-segmentation pipeline in Xenium Ranger 3.1.1. These improvements were visually confirmed in Xenium Explorer v3.2.0 by toggling between the original and imported segmentation layers. After this, the cell ID of all neurons containing more than two transcripts of all the identifier genes for each neuronal subcluster (together with RBFOX3 to exclude any glial cells) were grouped together and visualized using Xenium Explorer v3.2.0. Each cluster was visualized along with the boundary immunolabelling described above, and each side of the spinal cord exported as a PNG. These images were overlaid onto a laminar atlas, created using Grossman et al (2022) human lumbar L3 spinal cord atlas as a reference, in Adobe Photoshop 2025. Each cell was drawn in space using a single dot of the pen tool. The left and right sides of the spinal cord were individually rotated and resized to ensure the laminar boundaries were in the most accurate position and to be able to compare different shapes and sizes of spinal cord.

### Non-neuronal analysis

Neurons, astrocytes, microglia, oligodendrocytes, ependymal and endothelial cell sub-clustering was performed on individual Xenium spatial transcriptomic samples (n = 8), using Seurat (v4.3.0) in RStudio. Following initial quality control and cell-type annotation, each cell type was subset from each sample and analyzed separately. Gene expression data were scaled, and dimensionality reduction was performed using principal component analysis (PCA). The number of principal components used for downstream analysis was selected based on ElbowPlot inspection, retaining the top 10 components. UMAP was used for visualization, and clusters were identified using the Louvain algorithm (FindClusters, resolution = 1). Manual inspection and marker-based filtering were applied to remove doublets based on abnormal co-expression patterns.

To identify differentially expressed genes defining each cluster, we used FindAllMarkers function. Only genes expressed in at least 25% of cells within a cluster and showing a minimum log2 fold change of 0.25 were considered. P-values were adjusted for multiple testing using the Benjamini-Hochberg method to control the false discovery rate. Top marker genes (ranked by average log2 fold change) were grouped by cluster and were then annotated by putative biological function: homeostatic, vascular-associated, metabolic, or reactive, based on canonical astrocyte markers and transcriptional profiles. Due to the targeted nature of the Xenium panel and its limited gene coverage, functional classification was more constrained than in the single-nucleus dataset. However, this approach allowed for reliable classification of astrocyte, microglia, endothelial and ependymal subtypes and their spatial localization relative to neuronal populations of interest. We did also used the CCT label transfer and integration with the single nuclei sequencing subclusters to verify our classification was correct.

### Integration of human and mouse datasets

Cross species and rostro-caudal anatomy analysis of single-cell data was performed with the Seurat (v5.3.0) package in RStudio (v4.3.2). Briefly, human lumbar single-nucleus RNA-seq data was integrated with human cervical single-nucleus RNA-seq data using CCA integration. As 10x Genomics FLEX chemistry profiles only part of the transcriptome, we restricted analysis to the shared 18,135 genes available in both datasets. Joint feature set for integration anchors was identified as the intersection of top 5000 most variable genes in both datasets identified by the vst algorithm implemented in Seurat resulting in 2714 genes. After identifying integration anchors we performed CCA based data integration extending this to the full 18,135 gene set resulting in an adjusted gene-cell matrix. We used the latter as input to mouse-human dataset integration to overcome alignment problems stemming from single-cell chemistry (gene-specific-primer based FLEX chemistry for human lumbar vs. poly-A capture for cervical spinal cord) and species differences. In order to integrate mouse and human data, we first identified shared homologous genes using the mouse2human() function from the homologene R package converting all mouse genes to their human ortholog naming scheme. As a result, we excluded all mouse genes with no clear human ortholog as well as mouse genes with many corresponding human genes and excluded all human and mouse genes with ambiguous orthologs. The remaining merged dataset had 15,003 common genes that were retained for downstream analysis. Mouse and human data integration anchors were identified from an intersect of top 7500 most variable features between human and mouse data yielding 3479 genes for anchor point identification. Data integration was used to yield adjusted gene expression estimates for the full remaining 15,003 genes. We performed Leiden clustering on the resulting integrated gene-cell matrix with 30 PCs and clustering resolution=2. Cholinergic clusters (CHAT+ cells and associated two clusters) were removed from the final dataset and data reclustered yielding dataset with 50 transcripomic GABAergic and glutamatergic neuron types. Cells were named after the parent human lumbar neuron type if more than half of human lumbar neurons in that cluster belonged to a previously defined neuron class. If no majority lumbar human neuron type dominated the specific cluster, the neuron type was given a placeholder name. For cell-type prevalence analysis, we sampled the compared datasets with equal depth (mouse lumbar spinal cord vs. human lumbar spinal cord and human cervical spinal cord vs. human lumbar spinal cord). Bias in cell-type prevalence across compared samples was estimated by calculating the sum of corresponding neuron classes between the two samples and estimating % proportion belonging to each compared sample. Any deviation from 50% would be indicative of altered neuron-type prevalence.

Finally, to evaluate the conservation of cell-surface protein expression specificity across orthologous neuron types, we identified top differentially expressed cell surface protein encoding genes by aggregating the top 200 most variably expressed cell surface genes from each dataset (“vst” algorithm using the FindVariableFeatures() function in Seurat) with cell-surface protein encoding gene list defined by SurfaceGenie(*125*). This yielded a list of 315 most variable extracellular membrane associated protein encoding genes. We calculated the normalized pseudobulk expression (cell-type aggregated counts / # cells in type) for each of the genes in 49 of the orthologous neuron classes in each dataset and used spearman correlation to estimate the similarity of gene expression across neuron types in human lumbar, human cervical and mouse lumbar samples.

### Human spinal cord slice preparations for electrophysiological and morphological characterization

Human spinal cord tissue (∼1 cm in length; 3 - 4 pieces per donor) was obtained in the operating room and immediately placed in sealed 50-mL tubes containing NMDG-based solution pre-equilibrated with 95% O / 5% CO . The NMDG-based solution contained (in mM): 93 NMDG, 2.5 KCl, 1.25 NaH PO, 30 NaHCO, 20 HEPES, 25 Glucose, 5 Ascorbic acid, 2 Thiourea, 3 Sodium pyruvate, 10 MgSO, 0.5 CaCl, and 12 N-acetylcysteine; pH was adjusted to 7.3 with HCl, and osmolarity was 300 - 310 mOsm. Tissue was transported to the laboratory on ice. Upon arrival, spinal cord pieces were transferred to an ice-cold chamber containing NMDG-based solution continuously bubbled with 95% O / 5% CO and gently rinsed to remove residual blood. The tissue was then moved to a second ice-cold chamber with fresh NMDG-based solution continuously bubbled with 95% O / 5% CO, where the dura mater (when present) and dorsal and ventral rootlets were carefully trimmed using scissors and forceps, while leaving the menignes and pia matte intact. Spinal cords were glued onto a mounting cylinder by attaching the lateral surface to the cylinder, and molten 2% agarose was poured over the tissue until completely submerged. The agarose was rapidly condensed using a cooling block for 5 - 10 s. The agarose-embedded spinal cord block was inserted into the receiver of a Compresstome VF-200 (Precisionary Instruments) slicing platform. The slicing chamber was filled with carbogenated NMDG-based solution, and parasagittal slices (300 μm) were obtained for electrophysiological recordings. After sectioning, slices were transferred to pre-warmed (32 - 34 °C) carbogenated NMDG-based solution for 10 min. Following this initial recovery period, slices were transferred to a holding chamber containing room-temperature HEPES holding ACSF under continuous carbogenation. HEPES holding ACSF contained (in mM): 92 NaCl, 2.5 KCl, 1.25 NaH PO, 30 NaHCO, 20 HEPES, 25 Glucose, 5 Ascorbic acid, 2 Thiourea, 3 Sodium pyruvate, 2 MgCl, 2 CaCl ; osmolarity was 300 - 310 mOsm, and pH was ∼7.3 after carbogenation. The Compresstome VF-200-based slicing approach was adapted from the method previously described by the Feng laboratory for brain tissue (*126*) with minor modifications.

### Whole-cell patch-clamp recordings

Slices were transferred to a recording chamber containing ACSF continuously bubbled with 95% O / 5% CO for whole-cell patch-clamp recordings. ACSF was perfused at 2.0 mL/min using a Mini-Peristaltic Pump II (Harvard Apparatus). The recording ACSF contained (in mM): 124 NaCl, 2.5 KCl, 1.25 NaH PO, 24 NaHCO, 5 HEPES, 12.5 Glucose, 1 MgCl, and 2 CaCl ; osmolarity was 300 - 310 mOsm, and pH was ∼7.3 after carbogenation. Lamina II was identified as a translucent band across the dorsal horn under 4 x magnification on an upright microscope (BX50WI, Olympus). Whole-cell current-clamp recordings were then obtained from neurons in Lamina II under 40 x magnification using glass pipettes (4.0 - 5.0 MΩ) filled with a K-gluconate-based intracellular solution containing (in mM): 120 K-gluconate, 5 NaCl, 3 MgCl, 0.1 CaCl, 10 HEPES, 1.1 EGTA, 4 Na ATP, 0.4 Na GTP, and 15 Na - phosphocreatine. The intracellular solution was adjusted to pH 7.3 with KOH/HCl and to 290 mOsm with sucrose. For morphological reconstructions, biocytin (∼4 mg/ml) was added to the intracellular solution. Signals were acquired with a patch-clamp amplifier (Multiclamp 700B amplifier, Molecular Devices) and acquisition software (pClamp 11, Molecular Devices). Signals were low-pass filtered at 2 kHz and digitized at 10 kHz. No correction for the liquid junction potential was made. The experiments were conducted at room temperature.

After achieving a stable whole-cell configuration, resting membrane potential (RMP) and spontaneous activity were measured using a gap-free current-clamp recording without current injection for 1 min. Other intrinsic membrane properties and firing patterns were assessed in current-clamp mode using step and ramp current injections at the resting membrane potential or with the membrane potential held at −70 mV. For step injections, 1-s currents were delivered starting at −100 pA and increased in 25-pA increments until action potentials were elicited in three consecutive sweeps. This allowed determination of input resistance (Rinput), rheobase, action potential (AP) onset, AP voltage threshold (Vthreshold), AP amplitude, fast afterhyperpolarization (fAHP), voltage sag (Vsag), action potential duration at 50% AP amplitude (APD50), and AP kinetics (upstroke and downstroke velocities). Firing patterns and input-output relationships were then assessed using 1-s currents at 1x, 2x, 3x, and 4x rheobase. For ramp injections, 1-s depolarizing ramps starting from 0 pA and increasing in 25-pA increments were used to determine ramp rheobase; subsequent 1-s injections at 1x – 4x ramp rheobase were applied to characterize firing patterns and input-output relationships. After completion of whole cell current clamp recordings, the pipette was slowly withdrawn while applying gentle positive pressure to allow the ruptured membrane to reseal and re establish a gigaohm seal. Following electrode removal, recorded spinal cord slices were transferred to a continuously oxygenated ACSF chamber and incubated for 1 hour to allow intracellular biocytin diffusion throughout the soma and neuronal processes. Slices were then fixed overnight in 4% paraformaldehyde (PFA) at 4 °C. After fixation, slices were rinsed using 1× PBS and transferred to 1× PBS containing 0.05% sodium azide for long term storage prior to histological processing.

### Neuronal morphology and confocal imaging

The methods for neuronal morphological reconstruction and analysis were adapted from previously published studies (*127, 128*) with minor modifications, as briefly described below. Slices were rinsed in 1× Tris-buffered saline (TBS) for 5 min, seven times, and then incubated in 3% TBS-Triton for 1 h. Next, slices were incubated in 10% normal goat serum (NGS) with 0.5% bovine serum albumin (BSA) in TBS for 30 min, followed by two 5-min rinses in 1× TBS. Slices were then transferred to Streptavidin Alexa Fluor 488 conjugate (1:200) in TBS containing 1% NGS and 0.5% BSA and incubated for ∼16 h at 4 °C. On the following day, slices were rinsed four times for 5 min each in TBS and mounted on glass microscope slides using Dako Fluorescent mounting medium. Cover-slip shards (2 × 0.15 mm) were placed on either side of the slices to create a well and prevent compression, after which a thin cover glass was placed over the slices and shards, and the edges were sealed with nail polish. Confocal fluorescent images of recorded neurons were acquired using a Nikon Eclipse Ti inverted microscope equipped with four lasers (405, 488, 561, and 640 nm). Slices were scanned from top to bottom at 2-μm intervals using a 10× or 20× Plan Apo λ objective, and image acquisition was performed with NIS-Elements software (version 5.02).

### Data analysis

Input resistance was determined from the steady-state voltage responses to a series of 1-s hyperpolarizing current steps (−100 to 0 pA, 25-pA increments) as the slope of a linear least-squares fit to the resulting current-voltage relationship. Rheobase was defined as the smallest current step that elicited an action potential (AP). Voltage threshold for APs (in mV) was defined as the membrane potential at which dV/dt exceeded 10% of its maximum, relative to a dV/dt baseline measured 2 ms before the AP peak. AP amplitude was calculated as the difference between Voltage threshold and peak membrane potentials. AP half-width was measured at half-amplitude, using the first evoked AP for all measurements. AP onset was defined as the interval (in milliseconds) between current step onset and the voltage threshold of the first AP. Fast after-hyperpolarization (fAHP) following the first AP was measured as the difference between the AP threshold and the minimum membrane potential attained during the fAHP. Voltage sag (Vsag) was measured during a 1-s hyperpolarizing current step (−100 pA), and percent sag was calculated as 100 x (ΔVpeak - ΔVss)/ΔVpeak, where ΔVpeak is the peak voltage change and ΔVss is the steady-state voltage change. Electrophysiology data were analyzed using Clampfit (Version 11.2, Molecular Devices) and Prism version 9.2 GraphPad Software. Neuronal morphology was reconstructed, and soma size and neurite length were measured using ImageJ/Fiji software. Sholl analysis of biocytin-filled neurons was performed using a customized Simple Neurite Tracer plugin for Fiji (*127*). All figures were created using GraphPad Prism 9, Adobe Illustrator 2021, Adobe Photoshop 2021.

### Spinal cord slice preparation for S-DHPG recordings

Lumbar sections of the spinal cord were recovered from 2 organ donors (Transplant Quebec; a male aged and a female aged 58). The methods for preparing spinal cord slices were adapted from those described previously for non-human primate (*129*). The spinal cord was immersed in ice cold (0–4 °C) sucrose-based artificial cerebrospinal fluid (aCSF) of the following composition (mM): 100 sucrose, 63 NaCl, 2.5 KCl, 1.2 NaH2PO4, 1.2 MgCl2, 25 D-glucose and 25 NaHCO3, equilibrated with 95% O2-5% CO2. Parasagittal sections (350-550 μm) were cut using a vibratome (Leica VT1000s). Slices were subsequently incubated at 35 °C in NMDG-based recovery aCSF composed of the following composition (mM): 93 NMDG, 2.5 KCl, 1.2 NaH2PO4, 30 NaHCO3, 20 HEPES, 25 D-glucose, 5 sodium ascorbate, 2 thiourea, 3 sodium pyruvate, 10 MgSO4, 0.5 CaCl2, pH adjusted to 7.4 with HCl and equilibrated with 95% O2-5% CO2. Slices were then transferred to a recording aCSF of the following composition (mM): NaCl, 127; KCl, 1.9; KH2PO4, 1.2; CaCl2, 2.4; MgCl2, 1.3; NaHCO3, 26; D-glucose, 10; equilibrated with 95% O2-5% CO2 at 35 °C before cooling and maintenance of slices at room temperature prior to electrophysiological recording.

### Electrophysiological recordings with S-DHPG

Slices were transferred to a custom-built recording chamber and continuously perfused with aCSF at a rate of 6 ml.min-1 at a temperature of 35 ± 1°C. Whole-cell recordings were obtained from neurones in the translucent substantia gelatinosa layer of the superficial dorsal horn of spinal cord slices using a Multiclamp 700B amplifier or Axopatch 1D amplifier (Molecular Devices, California). Patch pipettes were pulled from thin-walled borosilicate glass (GC150TF-10; Harvard Apparatus) using a horizontal electrode puller (P-97 Flaming/Brown type micropipette puller; Sutter Instruments) and had resistances of between 6 and 12 MΩ when filled with intracellular solution of the following composition (mM): potassium gluconate, 140, KCl, 10, EGTA-Na, 1, HEPES, 10, Na2ATP, 4, Na-GTP, 0.3. Biocytin (5 or 10mM) was included in the solution to facilitate post-recording visualisation of neuronal morphology. Data were digitized at 10 kHz and stored on a PC running pCLAMP 10 data acquisition software (Molecular Devices, California). Recordings were primarily carried out in the current-clamp mode. However, spontaneous excitatory post-synaptic currents (EPSCs) and inhibitory post-synaptic currents (IPSCs) were recorded in the voltage-clamp configuration at holding potentials of −70 and −40mV, respectively. S-DHPG (Tocris Bioscience) was made up as a concentrated stock in DMSO, aliquoted and stored frozen until the day of experiments. Test compounds were diluted to required concentrations in aCSF and applied to the slices by superfusion at 35 ± 1°C from 50ml syringes, arranged in series with the main perfusion line. Upon completion of electrophysiological recording, slices were fixed in 4% paraformaldehyde in 0.1 M phosphate buffer (PB) composed of (M): 0.019 NaH2PO4·2H2O and 0.081 Na2HPO4, pH 7.4, before being washed in 0.1 M PB in preparation for subsequent immunohistochemistry. All analysis of electrophysiological data was undertaken using Clampfit 10 software (Molecular Devices, California). Results are expressed as mean + sem and statistical analysis performed using Student’s t-test (paired or unpaired as appropriate; Microsfoft Excel). Data was collected from 3 individual donors (22-year-old male, a 71 year old male and a 58 year old female.)

## Acknowledgements

The authors thank the organ donors and their families for their enduring gift, as well as our partners at Southwest Transplant Alliance and MidAmerica Transplant, without whom this research would not be possible. We also thank John Lemen and Dr. Gary Marklin of Mid-America Transplant for their invaluable support and assistance with tissue recovery. The authors also thank the Genome Center at The University of Texas at Dallas for the services to support our research. The authors are grateful to the Moody Flow Cytometry Facility at University of Texas Southwestern Medical Center for assistance with cell sorting. We gratefully acknowledge the members of the UTD recovery team: Keerthana Natarajan, Sera Nakisli, Joseph Lesnak, Moeno Kume, Lucy He, Marisol Mancilla Moreno, Urzula Enzastiga, Asta Arendt-Tranholm, Jayden O’Brien, Khadijah Mazhar, Kathleen Domalogdog, Ryan McKee, Nishka Kuttanna, Zulmary Manjarres Farias, Alejandro Otero Pedraza, Pegah Haghighi, Andi Wangzhou; and the recovery team at Washington University: Juliet M. Mwirigi, John Del Rosario, Maria Payne, Adam Dourson, and Grace Moore, for their invaluable contributions to this work.

## Author contributions

K.A.G & S.M.P performed single-nucleus sequencing of human spinal cord. K.A.G, O.C.D, S.M.P & J.M.B performed Xenium spatial sequencing. K.A.G, O.C.D, S.M.P, J.M.B, I.K & N.S performed computational analysis of the sequencing data. N.N.I, H.M, E.M, D.T-F optimized and performed reintegration of cell segmentations into the Xenium output data. S.I. optimized single-nucleus suspension generation and flow-assisted enrichment of neuronal nuclei. H.P. performed comparative mouse and human spinal cord analysis. A-H.P & S.I performed single-nucleus sequencing of mouse spinal cord. S.S., M.S.Y., E.V, P.H, T.K, A.C, J.C.R organized and performed tissue retrieval from human organ donors. L.S.M, R.S, J.L, P.G, R.A.S, R.T, R.J.H, B.A.C, D.S, R.G all contributed to the electrophysiological recordings. K.A.G, O.C.D, S.M.P, B.A.C, D.S, R.G, A-H.P & T.J.P wrote the manuscript. All authors edited the manuscript. K.A.G, O.C.D, A-H.P. & T.J.P devised the experimental plan. T.J.P, S.S., M.S.Y., & A-H.P. received funding to support the experimental work performed. A.P., G.D. A-H.P. & T,J.P. supervised the project.

## Declaration of Interests

Conflict of Interest Statement: T.J.P. is a co-founder of and holds equity in 4E Therapeutics, NuvoNuro, PARMedics, and Nerveli. T.J.P. has received research grants from AbbVie, Eli Lilly, Grunenthal, Evommune, Hoba Therapeutics, and The National Institutes of Health. The authors declare no conflicts of interest related to this work.

## Data and materials availability

Data and code used in this manuscript will be made publicly available. DOI for the human single nucleus sequencing data on SPARC.science is 10.26275/aikc-g5tv and for the Xenium data is 10.26275/7ozi-u6bk. Code for cross-species comparative analysis is available at https://github.com/PoolLab/human_spinal_cord.

## Ethics approval

Human tissue procurement procedures were approved by the Institutional Review Board at the University of Texas at Dallas (MR-15-237). The Southwest Transplant Alliance (STA) obtained informed consent for research tissue donation from first-person consent or from the donor’s legal next of kin. Policies for donor screening and consent are those established by the United Network for Organ Sharing (UNOS). STA follows the standards and procedures established by the US Centers for Disease Control (CDC) and are inspected biannually by the Department of Health and Human Services (DHHS). The distribution of donor medical information is in compliance with HIPAA regulations to protect donor privacy. Post-mortem human tissue procurement at Washington University in St Louis (WashU) was approved by the Human Research Protection Office (IRB exemption 202203041) in collaboration with Mid-America Transplant.

## Funding Statement

This research was supported by the National Institute Of Neurological Disorders and Stroke of the National Institutes of Health through the PRECISION Human Pain Network (RRID:SCR_025458), part of the NIH HEAL Initiative (https://heal.nih.gov/) under award number U19NS130607 to RWG and U19NS130608 to TJP. A.H.P. is supported by Eugene McDermott Endowed Scholars Program, Rita Allen Foundation Award in Pain and McKnight NBD Award. The content is solely the responsibility of the authors and does not necessarily represent the official views of the National Institutes of Health.

## Notes

### Summary of Updates

We added additional analysis on RNA sequencing data and added electrophysiology data on human spinal cord. Authors were added to the paper to account for the new data and analysis that was added.

