## Supplemental figures and tables for "A molecular map of the human spinal dorsal and ventral horn defines arrangement of neuronal types and glial sex differences"

| **Donor #** | **Age** | **Sex** | **COD** | **# Cells**  **DH VH** | | **Median reads per nuclei**  **DH VH** | | **Median genes per nuclei**  **DH VH** | | **Median UMI counts per nuclei**  **DH VH** | | **Total genes detected**  **DH VH** | |
| --- | --- | --- | --- | --- | --- | --- | --- | --- | --- | --- | --- | --- | --- |
| 1 | 23 | Female | Anoxia | 6,179 | 6,845 | 6,448 | 6,775 | 2,009 | 2,069 | 3,207 | 3,380 | 16,897 | 16,736 |
| 2 | 20 | Male | GSW | 4,935 | 7,729 | 7,637 | 7,166 | 2,299 | 2,220 | 3,813 | 3,568 | 17,451 | 17,159 |
| 3 | 32 | Male | GSW | 5,401 | 5,160 | 6,870 | 6,697 | 2,164 | 2,136 | 3,405 | 3,327 | 16,960 | 16,489 |
| 4 | 36 | Female | Anoxia OD | 6,225 | 6,992 | 5,829 | 4,824 | 1,872 | 1,574 | 2,896 | 2,399 | 16,744 | 16,576 |
| 5 | 43 | Male | MVA | 4,173 | 6,207 | 7,573 | 5,551 | 2,288 | 1,801 | 3,784 | 2,765 | 16,743 | 16,582 |
| 6 | 46 | Female | MVA | 5,594 | 5,734 | 6,368 | 6,098 | 1,977 | 1,900 | 3,179 | 3,027 | 17,041 | 16,658 |
| 7 | 50 | Female | Anoxia | 7,419 | 3,340 | 8,051 | 8,117 | 2,429 | 2,427 | 4,010 | 4,039 | 16,885 | 16,619 |
| 8 | 56 | Male | Anoxia | 5,653 | 5,307 | 5,138 | 6,446 | 1,746 | 2,079 | 2,548 | 3,198 | 17,176 | 17,169 |
| 9 | 39 | Male | Anoxia OD | 12,810 | 4,970 | 7,814 | 11,975 | 1,954 | 2,722 | 3,211 | 4,878 | 17,788 | 17,804 |
| 10 | 37 | Female | GSW | 14,319 | 2,341 | 13,967 | 17,409 | 3,285 | 3,506 | 5,707 | 7,100 | 17,720 | 16,688 |
| 11 | 60 | Female | Anoxia | 6,816 | 7,914 | 14,512 | 11,550 | 3,260 | 2,635 | 5,930 | 4,725 | 17,780 | 17,466 |

Supplemental Table 1. Quality control metrics per sample across all 11 donors divided by dorsal horn and ventral horns.

(Attached as separate excel file)

Supplemental Table 2. A list of genes included on the custom 480 gene 10X Xenium panel.

(Attached as separate excel file)

Supplemental Table 3. Pseudobulk Expression of Variable Cell Surface Genes in Mouse and Human Lumbar Spinal Cord

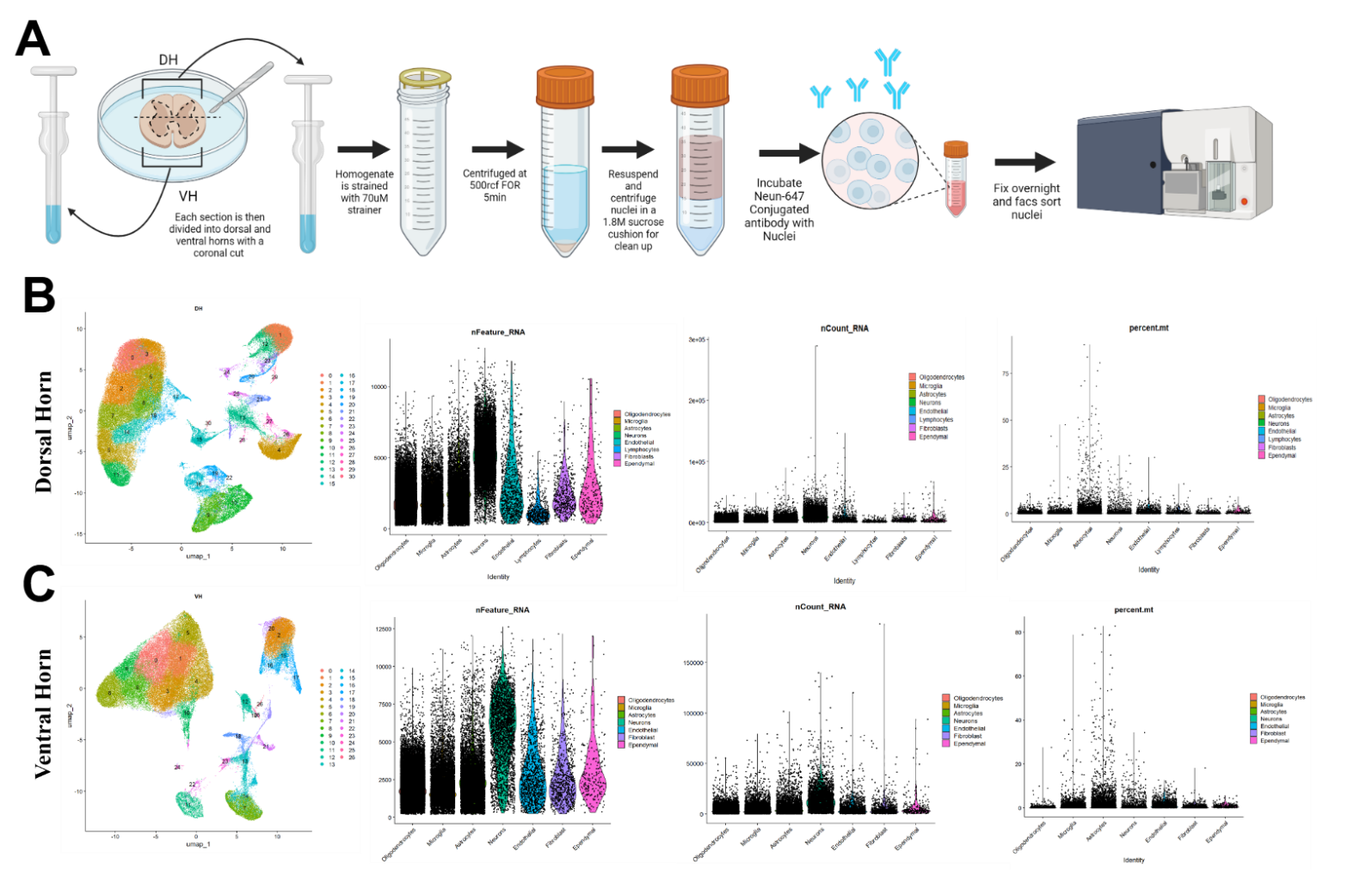

Supplemental Figure 1. (**A**) Workflow of single nuclei sequencing extraction in the human spinal cord using FACS sorting. (**B**) UMAP showing 31 clusters of the dorsal horn and the violin plots for three quality control metrics across all cells demonstrating high number of detected features, a good number of RNA molecules, and low percentage of mitochondrial gene expressions. (**C**) UMAP showing 30 clusters of the ventral horn and the violin plots for three quality control metrics across all cells demonstrating high number of detected features, the RNA molecules, and low percentage of mitochondrial gene expressions.

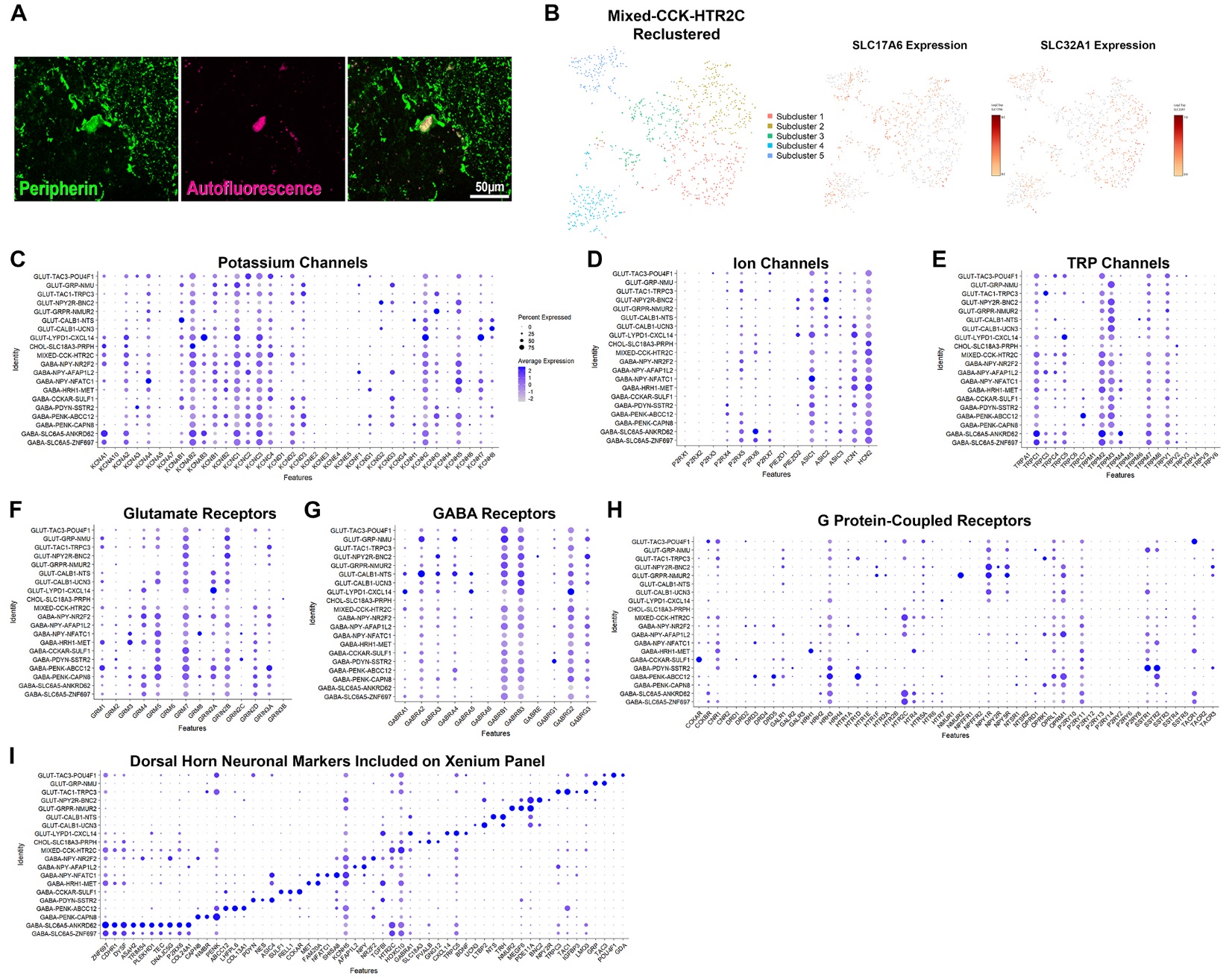
Supplemental Figure 2. (**A**) Peripherin-immunolabelling reveals positively labelled cell bodies in the dorsal horn of the human spinal cord. (**B**) Subclustering the mixed dorsal horn population of neurons (MIXED-CCK-HTR2C) revealed five transcriptomically similar clusters, all containing excitatory (SLC17A6+) and inhibitory (SLC32A1+) nuclei. Dot plots showing expression of potassium channel components (**C**), ion channels including purinurgic and ASIC channels (**D**), TRP channels (**E**), glutamate receptor subunits (**F**), GABA receptor subunits (**G**), G protein-coupled receptors (**H**) and marker genes used to identify each population on the Xenium panel (**I**) in dorsal horn neuronal clusters.

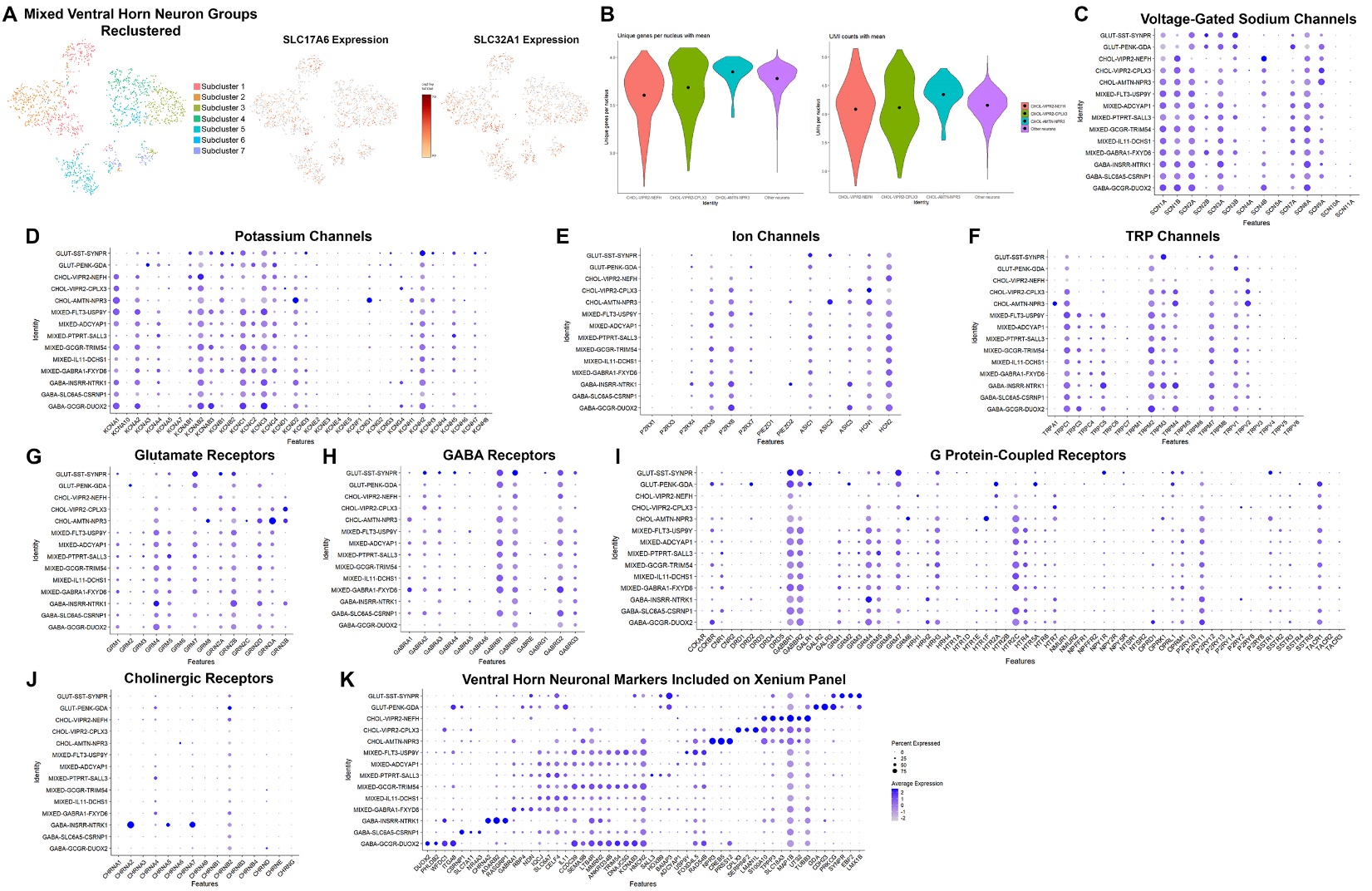
Supplemental Figure 3. **A**, Subclustering the six mixed ventral horn populations of neurons produced seven transcriptomically similar clusters, all containing excitatory (SLC17A6+) and inhibitory (SLC32A1+) nuclei. (**B**) Unique genes and UMIs per nucleus of the three motor neuron groups compared to other interneuron populations shows that the motor neurons clusters do not have a lower quality of nucleus in general. Dot plots showing expression of voltage-gated sodium channels (**C**), potassium channel components (**D**), ion channels including purinurgic and ASIC channels (**E**), TRP channels (**F**), glutamate receptor subunits (**G**), GABA receptor subunits (**H**), G protein-coupled receptors (**I**), cholinergic receptors (**J**), and marker genes used to identify each population on the Xenium panel (**K**) in ventral horn neuronal clusters.

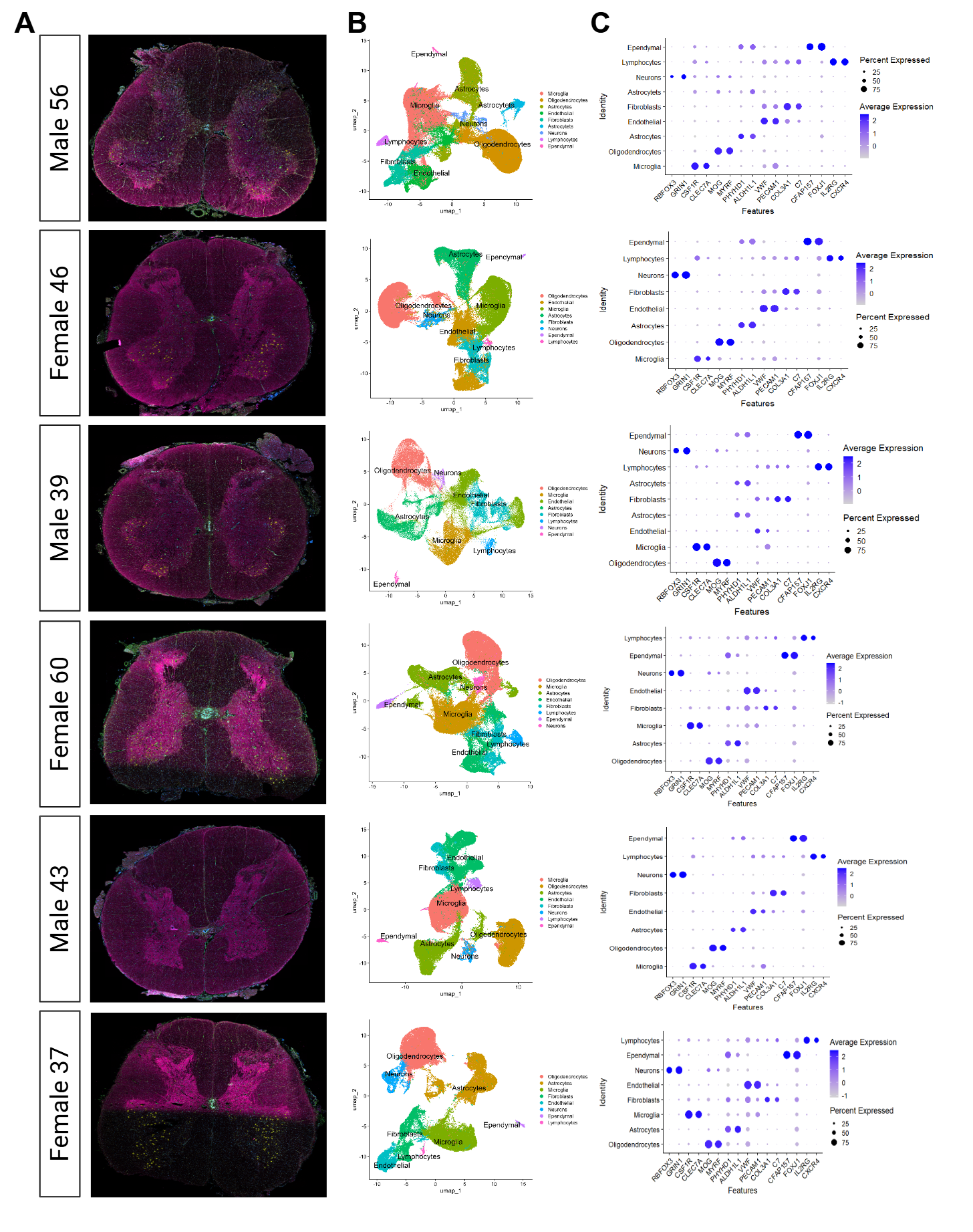

Supplemental Figure 4. Xenium spatial transcriptomics on each individual sample**.** (**A**) Initial cell segmentation output from Xenium using Xenium explorer across six samples. (**B**) UMAP plots showing named clusters for each of the six individual samples. (**C**) Dot plot displaying the marker genes used to annotate clusters across the six samples.

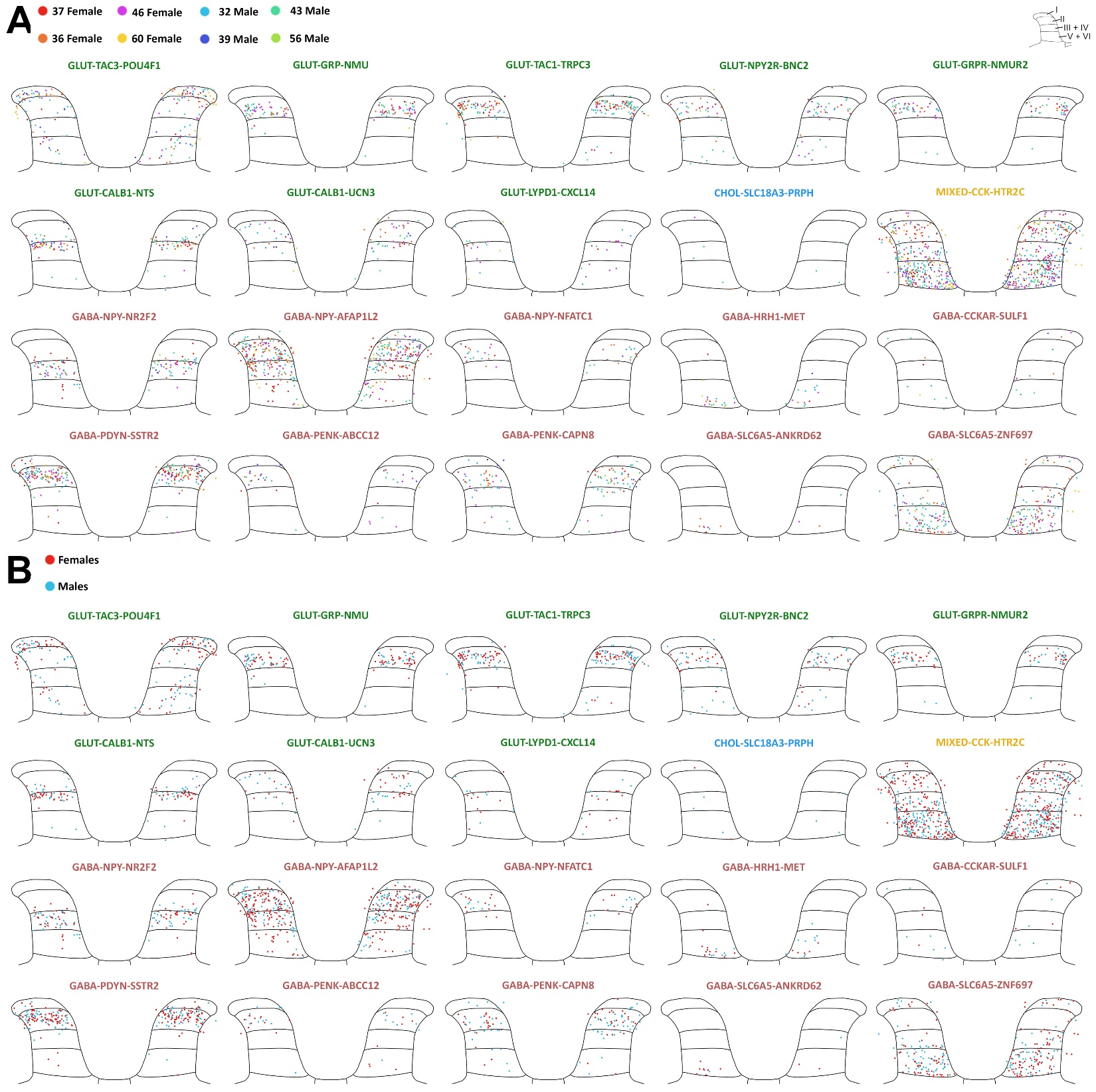

Supplemental Figure 5. Spatial organization of dorsal horn clusters shown per individual donor and grouped by sex.

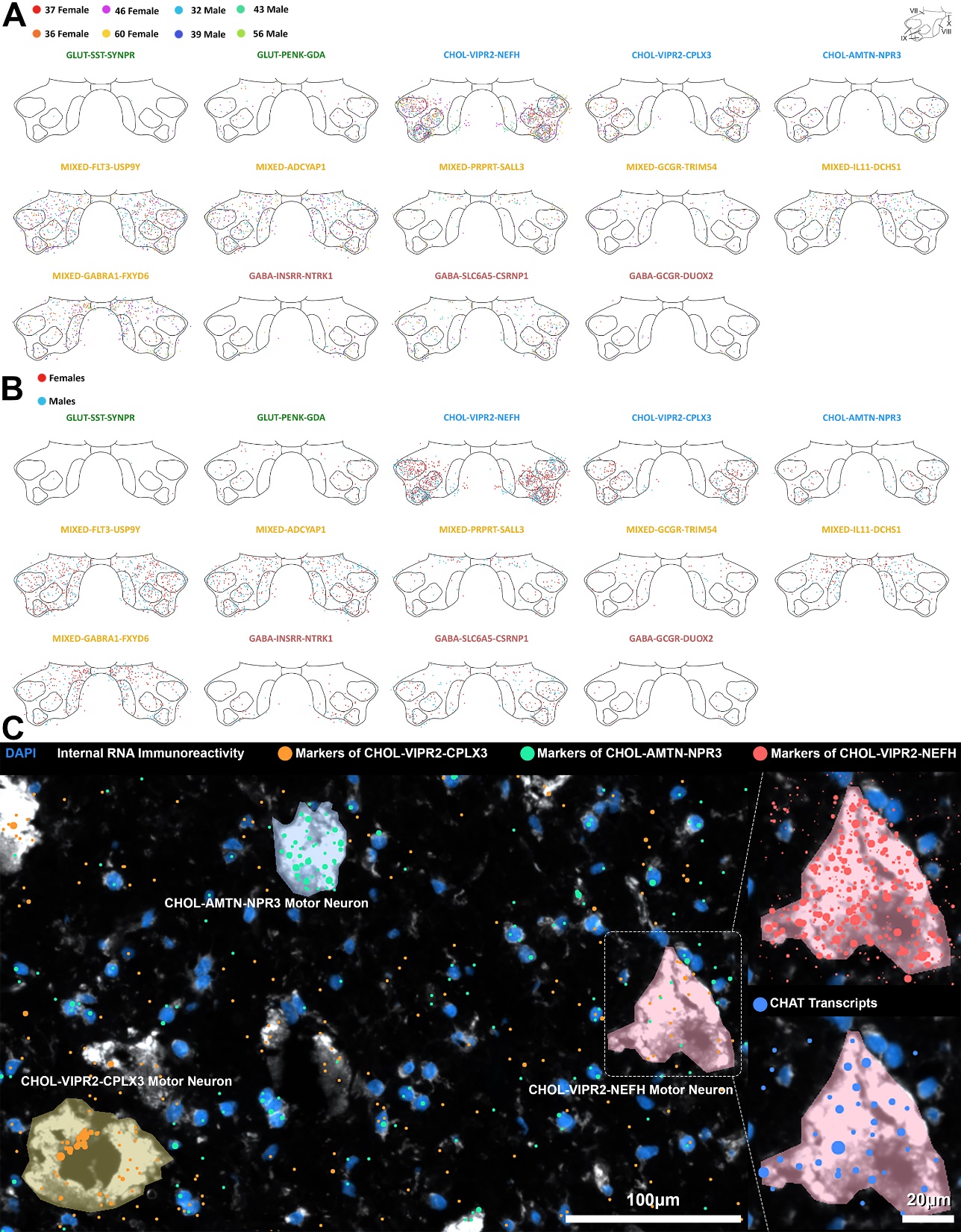

Supplemental Figure 6. Spatial organization of ventral horn clusters shown per individual donor (**A**) and grouped by sex **(B**). Image from Xenium Explorer showing DAPI and internal RNA immunostaining included in the Xenium cell segmentation module, detailing large cells assigned to each of the three motor neuron clusters: CHOL-VIPR2-NEFH (pink), CHOL-VIPR2-CPLX3 (yellow) and CHOL-AMTN-NPR3 (blue; **C**) based on transcript expression. The CHOL-VIPR2-NEFH motor neuron mostly lacks markers of CHOL-VIPR2-CPLX3 (CPLX3, SERPINF2 & LMAN1L orange transcripts) or CHOL-ATMN-CPLX3 (NPR3, CREB5 & PRSS12, blue transcripts), but is enriched in all CHOL-VIPR2-NEFH markers (S100A10, TPPP3, SLC18A3, MAP1B, UTS2, TUBB3, red transcripts, inset) and cholinergic transcripts such as CHAT (blue transcripts, inset).

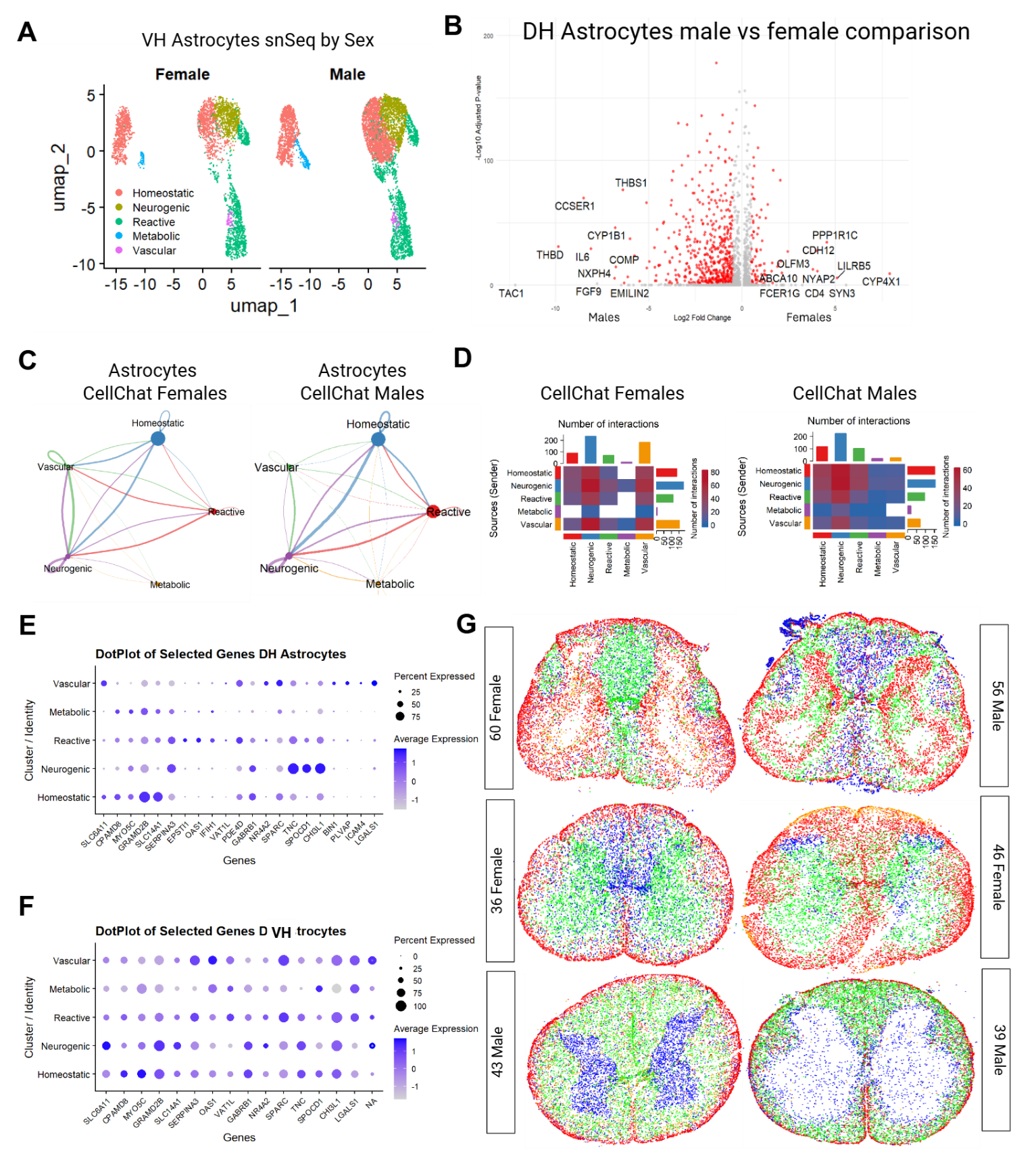

Supplemental Figure 7. (**A)** UMAP showing subclustered ventral horn astrocytes split by sex. (**B)** volcano plot demonstrating differential gene expression changes between males and female in dorsal horn astrocytes. (**C**). Intercellular communication network reflecting the numbers and intensity of interactions between the functional categories in the dorsal horn astrocytes split by males and females. (**D)** Representative heatmap of the intercellular communication network between astrocytes in males and females. (**E)** Dot plots demonstrating genes of interest for categorizing astrocytes by function in the dorsal horn. (**F**). Dot plots of categorizing genes in the ventral horn. (**G**) Spatial organization of astrocytes categorized by function across 6 remaining donors.

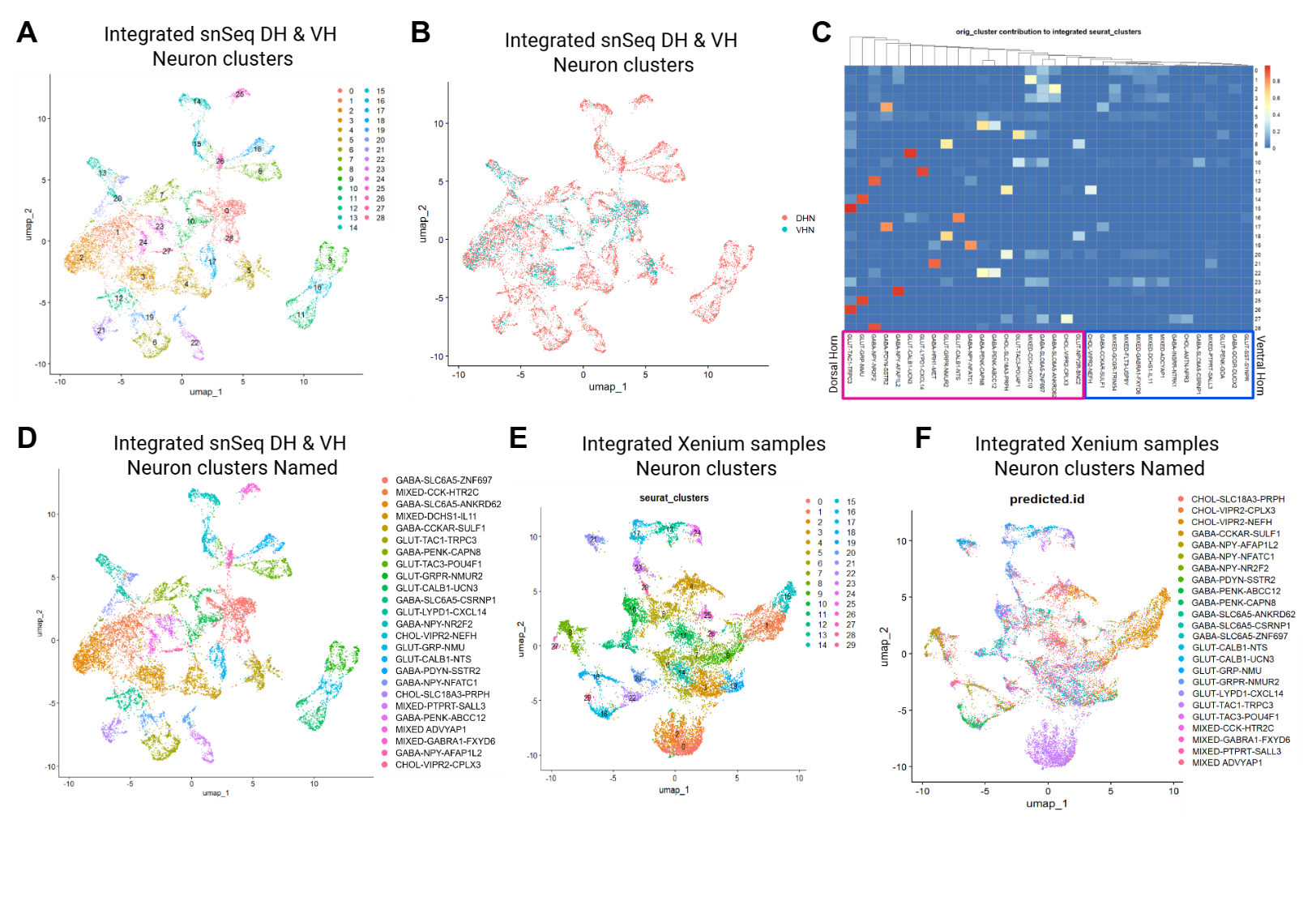

Supplemental Figure 8. (**A**) UMAP of integrated ventral horn and dorsal horn neuronal clusters yielding 29 clusters. (**B**) UMAP of integrated neuronal clusters showing distribution of ventral horn (blue) and dorsal horn (red) nuclei. (**C**) heatmap to visualize the outcome of label transfer, where predicted cell type annotations from the dorsal horn and ventral horn neuronal clusters were mapped onto the clusters of the integrated neuronal object. (**D**) UMAP depicting the final cluster assignment of the integrated single nuclei sequencing neuronal clusters. (**E**) UMAP of integrated xenium neuronal clusters. (**F**) UMAP of final cluster assignment of the integrated xenium neuronal clusters using label transfer.

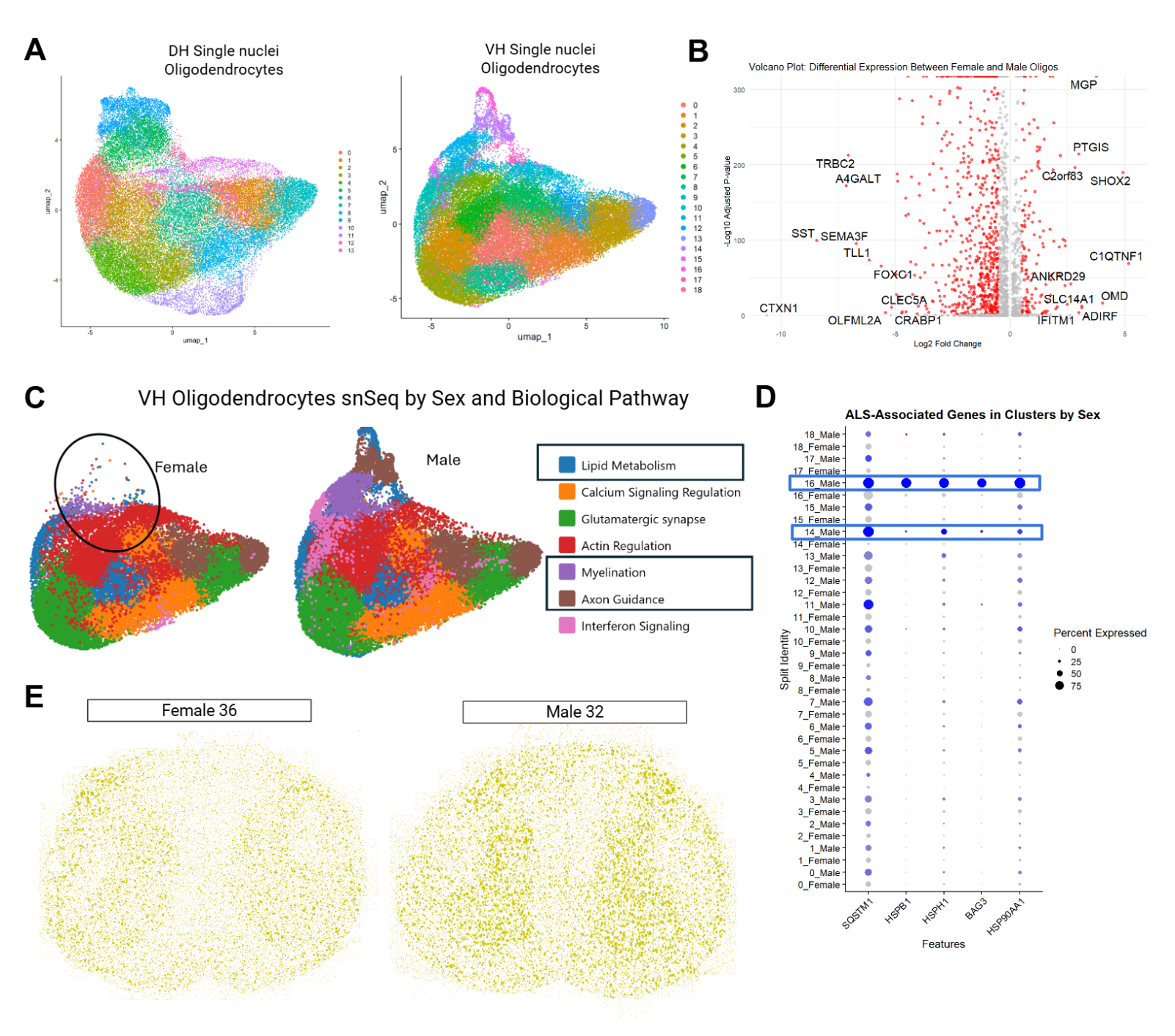

**Supplemental Figure 9**: (**A**) single nuclei sequencing UMAPs of oligodendrocytes from the dorsal horn (14 clusters) and ventral horn (19 clusters). (**B**) Volcano plot demonstrating differential gene expression changes between males (left) and female (right) in dorsal horn astrocytes. (**C**) single nuclei sequencing UMAPs of oligodendrocytes split by sex and categorized by biological pathway. (**D**) Dot plot looking at ALS associated genes that are highly expressed in clusters that are male exclusive (14 & 16). (**E**) Xenium spatial distribution of oligodendrocytes in two representative male and female donors.

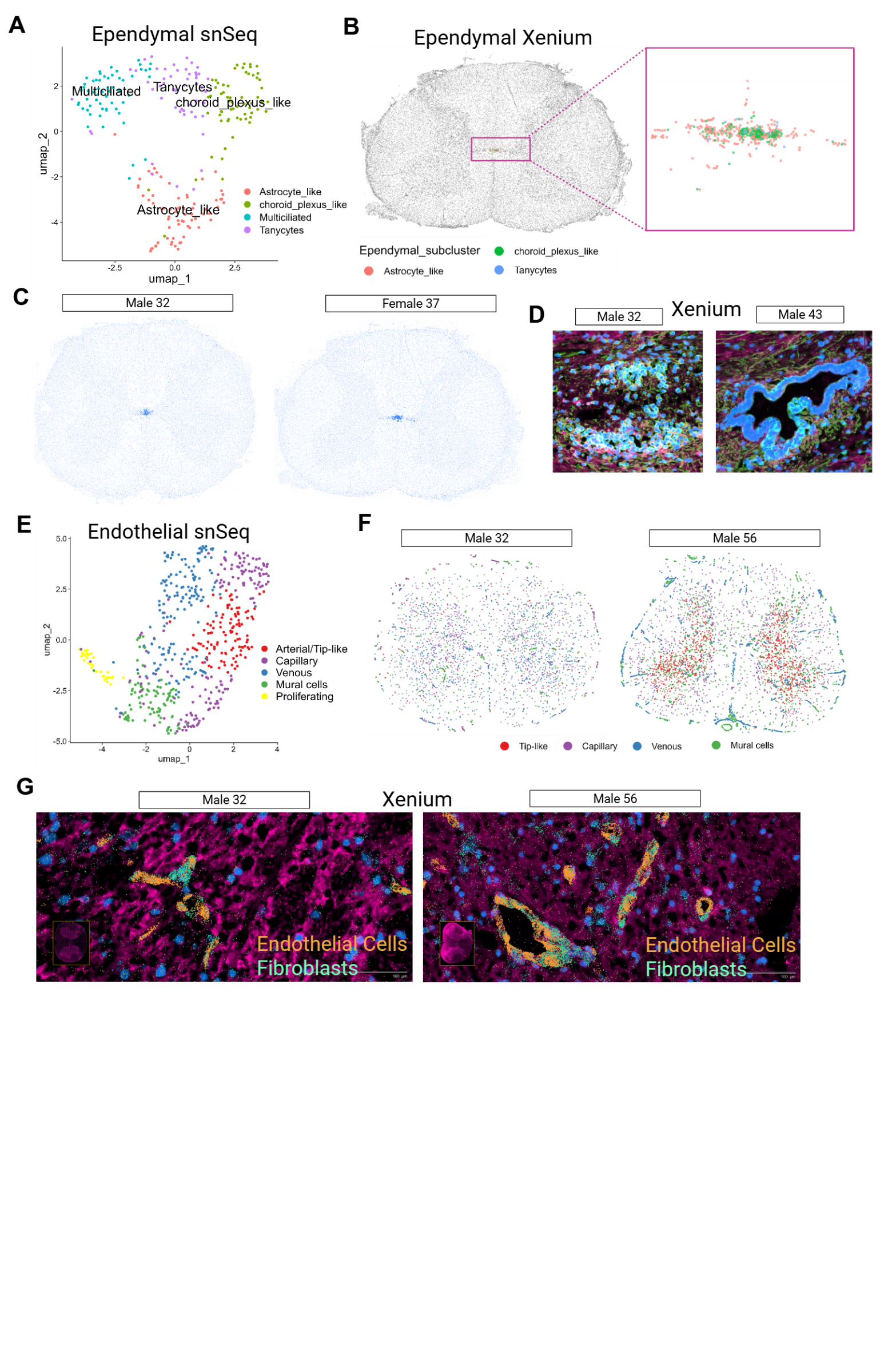

**Supplemental Figure 10**: (**A**) single nuclei sequencing UMAP of subclustered endothelial cell categories. (**B**) Spatial distribution of endothelial cells in one xenium sample (**C**) Representative male and female samples of endothelial cell in xenium. (**D**) Xenium image of differences in stenosis of central canal between two donors demonstrating it is age independent. (**E**) single nuclei sequencing UMAP of subclustered endothelial cell categories. (**F**) Xenium spatial distribution of endothelial cells between a younger and an older male sample. (**G**) Xenium image of differences in size of vascular formations across two different aged samples in the dorsal horn.

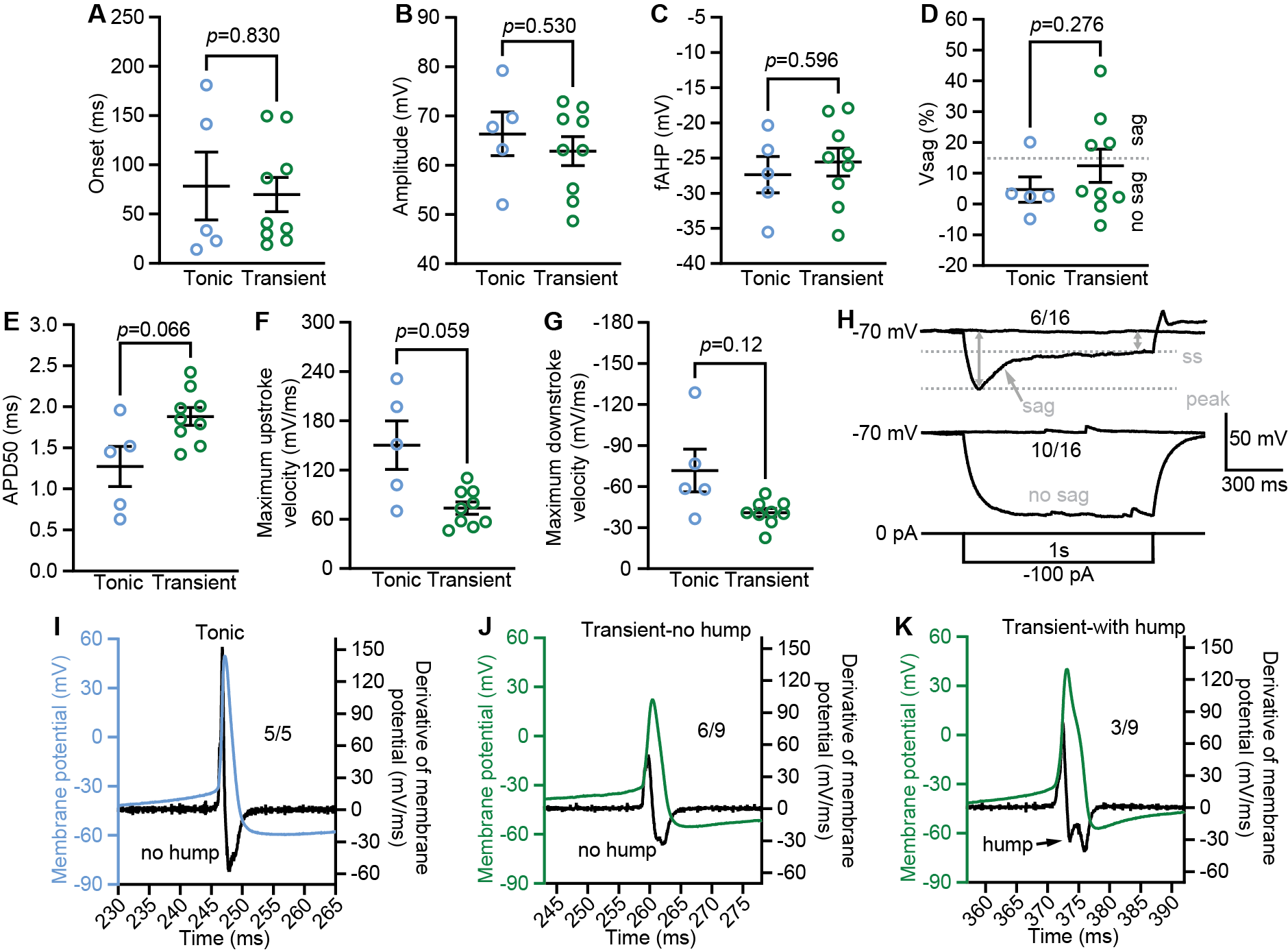

**Supplemental Figure 11:** Electrophysiological characterization of tonic and initial firing neurons in lamina II of the human spinal dorsal horn (hSC). (A-F) Comparison of intrinsic properties between tonic and initial firing hSC lamina II neurons, including action potential (AP) latency (onset, A), AP amplitude (B), fast after-hyperpolarization (fAHP, C), voltage sag (Vsag, D) and AP half-width (APD50, E). Two tailed unpaired t-test. (G, H) Comparison of maximum upstroke (G) and downstroke velocity (H) of APs between tonic and initial firing hSC lamina II neurons. Two tailed unpaired t-test. (I-K) Representative action potential traces and corresponding membrane potential derivatives from tonic (I) and initial (J, K) firing hSC lamina II neurons. A small subset of initial firing neurons exhibited a repolarization hump during the AP in the derivative trace (K, indicated by the arrow); however, this feature was not observed in the larger population of initial firing neurons (J) or in any of the recorded fast firing neurons (I). (L) Representative membrane potential traces in response to a -100 pA hyperpolarizing current injection (1 s duration) from a holding potential of -70 mV. Within the total recorded population (n=16), a subset of neurons displayed a prominent sag of the membrane potential (indicated by the grey arrow) to a steady-state (ss) level but the other population of neurons showed a stable hyperpolarized membrane potential without a visible sag.
